# Blocking antibodies against integrin-α3, -αM, and -αMβ2 de-differentiate myofibroblasts and reverse lung and kidney fibroses

**DOI:** 10.1101/2021.06.07.447405

**Authors:** Michael JV White, Melis Ozkan, Jorge Emiliano Gomez-Medellin, Michal M Rączy, Kyle M Koss, Ani Solanki, Zheng Jenny Zhang, Aaron T. Alpar, Bilal Naved, Jason Wertheim, Jeffrey A Hubbell

## Abstract

Fibrosis is involved in 45% of deaths in the United States, and no treatment exists to reverse the progression of the disease. Myofibroblasts are key to the progression and maintenance of fibrosis. We investigated features of cell adhesion necessary for monocytes to differentiate into myofibroblasts, seeking to identify pathways key to myofibroblast differentiation. Blocking antibodies against integrins α3, αM, and αMβ2 de-differentiate myofibroblasts *in vitro*, lower the pro-fibrotic secretome of myofibroblasts, and reverse lung and kidney fibrosis *in vivo*. Decorin’s collagen-binding peptide directs blocking antibodies (against integrins-α3, -αM, -αMβ2) to both fibrotic lungs and fibrotic kidneys, reducing the dose of antibody necessary to reverse fibrosis. This targeted immunotherapy blocking key integrins may be an effective therapeutic for the treatment and reversal of fibrosis.

**Summary:** Blocking antibodies against integrins-α3, -αM, and -αMβ2 can be targeted to sites of fibrosis, reverse lung and kidney fibroses, and offer the potential to bring immunotherapy to fibrosis

## Introduction

Fibrosis is defined by dysregulated extracellular matrix (ECM) deposition leading to scar tissue deposition and increases in tissue stiffness [1]. Fibrosing diseases—including pulmonary fibrosis, congestive heart failure, liver cirrhosis, and end-stage kidney disease—are involved in 45% of deaths in the United States [1, 2]. There are only 2 FDA approved treatments for fibrosis, pirfenidone and nintedanib [2, 3]. Pirfenidone and nintedanib slow, but do not reverse, the progression of fibrosis [4], with mechanisms of action that are poorly understood [5].

A major goal of research in fibrosis is developing a treatment capable of reversing established fibrosis [1]. To our knowledge, only one treatment (recombinant pentraxin-2, PRM-151) has thus far shown even a modest ability to reverse even some symptoms of fibrosis in some patients [6]. To fully reverse fibrosis, several cell and ECM-related changes must occur: myofibroblasts must de-activate to reduce the stiffness of the tissue, collagen deposition must cease, and existing ECM must be remodeled.

Interrupting collagen deposition alone can destabilize scar tissue [1] and monocyte-derived cells are capable of removing deposited ECM while regenerating tissue [7].

Myofibroblasts are cells that adopt a spindle-shaped morphology and actively increase surface stiffness and tension through exertion of force through their cytoskeleton’s attachment to the extracellular matrix. Myofibroblasts also secrete pro-fibrotic factors and collagen [1, 8–10]. Myofibroblasts are a heterogenous population that arise from multiple progenitor cells, and the term “myofibroblast” denotes the function of a cell rather than its origin [11]. Myofibroblasts can arise from several progenitors, including hepatocytes, fibroblasts, epithelial cells, and monocytes [1, 11, 12].

Myofibroblasts are not found in normal, healthy tissues, and are formed or recruited in response to injury [12, 13]. Myofibroblast de-activation is the deciding factor between tissue damage being resolved in regeneration or by continued scarring and fibrosis [9]. Myofibroblasts can de-activate through apoptosis [14] or de-differentiation [12, 15].

Uniquely among myofibroblast precursors, monocytes can be recruited from the bloodstream to different fibrotic tissues in the body [16]. These monocyte-derived myofibroblasts may provide a up to a third of the total number of myofibroblasts in liver [17], kidney [18], lung [19, 20], and skin [21] fibroses. Whether monocyte-derived cells assume an anti-fibrotic or a pro-fibrotic differentiation state can determine whether tissue damage is resolved with regeneration or progresses to fibrosis [22, 23].

Integrins are membrane glycoproteins that are involved in mediating interactions between cells and ECM surfaces [24]. Integrins comprise 18 α subunits and 8 β subunits, which combine to form 24 αβ heterodimers [25]. Each αβ heterodimer has a unique binding profile, which allows integrins a broad array of integrin ligands, including ECM proteins, soluble ligands, or other membrane proteins [25]. Integrins transduce signals from the ECM to the cytoskeleton in what is known as “outside-in” signaling, and transduce force from the cytoskeleton to the ECM in “inside-out” signaling [24].

Integrins can be activated by extracellular interactions that induce conformational changes to integrin’s intracellular domains. Conversely, integrins can be activated by intracellular cytoskeletal interactions that increase integrin’s ligand binding through conformational changes to integrin’s extracellular domains [24, 26]. Among the proteins capable of activating integrins are the intracellular tension-sensing proteins talin1 and talin2 [27]. Talins activate integrins through interactions with the β integrin tail [24], which induces a conformational change to the αβ integrin’s extracellular domain that increases affinity for ligands.

Focal adhesions (FAs) are subcellular structures rich in integrins which bind to the extracellular ECM, and intracellular talins which bind to the cytoskeleton [28–30]. FAs are dynamic, deformable, and responsive structures at the cell membrane that integrate signals and transmit/transduce force from the intracellular actin cytoskeleton, transmembrane integrins, and ECM [31–34]. Integrins interact with multiple ECM proteins [31, 32], and are essential to both cellular tension sensing, the formation of FAs, and the maintenance of FAs [35]. FAs are transient in migrating cells, but can become super-mature in myofibroblasts [29]. Super-mature FAs are both physically larger [29] and have increased concentrations of integrins, talins, and cytoskeletal proteins [28, 35].

Fibrillar adhesions (FBs) are complexes of cytoskeletal machinery and actin that connect to FAs and localize into long intracellular bundles throughout myofibroblasts [36]. Many proteins interact in these FAs and FBs, leading to a complex signaling and force-distribution environment [28].

The processes by which myofibroblasts respond to—and influence—their mechanical environment are called mechanosensing and mechanotransduction, respectively. Mechanosensing is a dynamic process integrating multiple signals between surface integrins, intracellular cytoskeleton, tension sensing talins, and secreted signals [8]. Changes in the interactions between integrins, intracellular proteins, and the actin cytoskeleton can lead to signaling changes in what might otherwise appear as a static cellular environment [31].

Integrin-αM is found on monocyte-derived cells [24]. αM forms an αβ heterodimer with integrin-β2, and β2 is also found on monocyte-derived cells [24]. αM and β2 together form the heterodimer αMβ2 (called MAC1). αMβ2 binds over 30 different ligands, including fibrinogen, and is also a canonical member of the complement pathway (CR3) [37, 38]. αMβ2-positive monocyte-derived cells are key to the development of fibrosis [39]. CRBM1/5 is a mouse anti-human blocking antibody that binds a conformational epitope found only on αMβ2, not on αM alone [40]. Leukadherin is a small molecule that promotes αMβ2 association [41].

Integrin-α3 is an integrin used to bind laminin [42]. α3 forms an αβ heterodimer with integrin-β1, and is found on both fibroblasts and monocytes.

Decorin’s heparin-binding domain contains a collagen-binding peptide (CBP) that has affinity for fibrotic tissues, and that decoration of antibody with CBP increases retention in fibrotic lungs [43].

Here we show that CBP-functionalized blocking antibodies against αMβ2 (CBP-α-αMβ2), αM (CBP-α- αM), and α3 (CBP-α-α3) de-differentiate myofibroblasts, lower the pro-fibrotic secretome of myofibroblasts, and reverse fibrosis in mouse lung and kidney models. These targets were generated by an mRNAseq study comparing monocytes cultured under anti- and pro-fibrotic conditions [44]. These results raise the possibility of using targeted anti-integrin antibodies as an immunotherapy against fibrosis.

## Results

We began this study with the observation that monocyte differentiation into myofibroblasts was dependent on adhesion to a stiff surface [44]. We sought to identify key pathways of monocyte-to-myofibroblast differentiation by performing an mRNAseq comparison of monocytes cultured on soft (1 kPa; not allowing myofibroblast differentiation) and stiff (12 kPa; allowing myofibroblast differentiation) surfaces (Table 1). We identified a key role for integrins α3, αM, and αMβ2, and blocking antibodies against these integrins de-differentiated existing myofibroblasts and reversed fibrosis in a murine model of pulmonary fibrosis.

**Table 1:** Changes in mRNA expression from culturing monocytes on soft, anti-fibrotic (1 kPa) and stiff, pro-fibrotic (12 kPa) surfaces, assessed by RNAseq. Enrichment score is calculated based on the maximum deviation from 0 in the pathway analysis [78]. In this analysis, 5.18 is the highest possible score.

Tables and Supplemental Figures
| Upregulated pathways | Functional annotation clustering | Enrichment score | Investigated protein targets | Fold change | Investigated protein targets | Statistical significance |
| --- | --- | --- | --- | --- | --- | --- |
| Chemokine signaling | Complement activation | 5.18 | ITGAM | 1.797671 | ITGB2 | 9.15E-04 |
| Leukocyte extravasation signaling | Chemokine signaling | 5.18 | ITGA3 | 3.536679 | ITGB1 | 2.59E-02 |
| IL-6 signaling | Phagocytosis recognition | 5.18 | ITGA7 | 4.487744 | ITGAX | 4.75E-05 |
| Paxillin signaling | positive regulation of inflammatory response | 5.18 |  |  |  |  |
| Integrin signaling | monocyte chemotaxis | 5.18 |  |  |  |  |
| IL-13 signaling | receptor-mediated endocytosis | 5.18 |  |  |  |  |

Our mRNAseq study of monocytes cultured on soft (1 kPa) and stiff (12 kPa) surfaces revealed that integrins αM (ITGAM), α3 (ITGA3), and α7 (ITGA7) were directly upregulated by culture on the stiff surface (Table 1). Upstream regulators of β1 (ITGB1), β2 (ITGB2), and αX (ITGAX) were also upregulated. Interestingly, no TGFβ-specific integrin or upstream regulator was found to be upregulated, including ITGAV, ITGA11, ITGB3, ITGB5, ITGB6, ITGB7 [8], nor was TGFβ itself. Additionally, several upregulated pathways and functional annotation clustering also indicated the involvement of integrins in the differentiation of myofibroblasts (Table 1).

To confirm the results of our mRNAseq study, we assessed the ability of anti-integrin antibodies to promote or inhibit myofibroblast differentiation. Blocking antibodies against α3 (α-α3), αM (α-αM), and against the heterodimer of αM and β2 (α-αMβ2) consistently de-differentiated monocyte- myofibroblasts (as determined by morphology) when added to culture (Figure 1A, IC50 in Table 2), while α-α7, α-β1, α-αX inconsistently de-differentiated monocyte-myofibroblasts, and α-β2 caused apoptosis (data not shown, [45]). An antibody that stabilizes the association of integrin α2β1 (clone Gi14) promoted myofibroblast differentiation (Figure 1A, [46]). The small molecule leukadherin, which increases the associated of αM and β2, also promoted myofibroblasts differentiation (Figure 1B, [41]).

**Figure 1:**
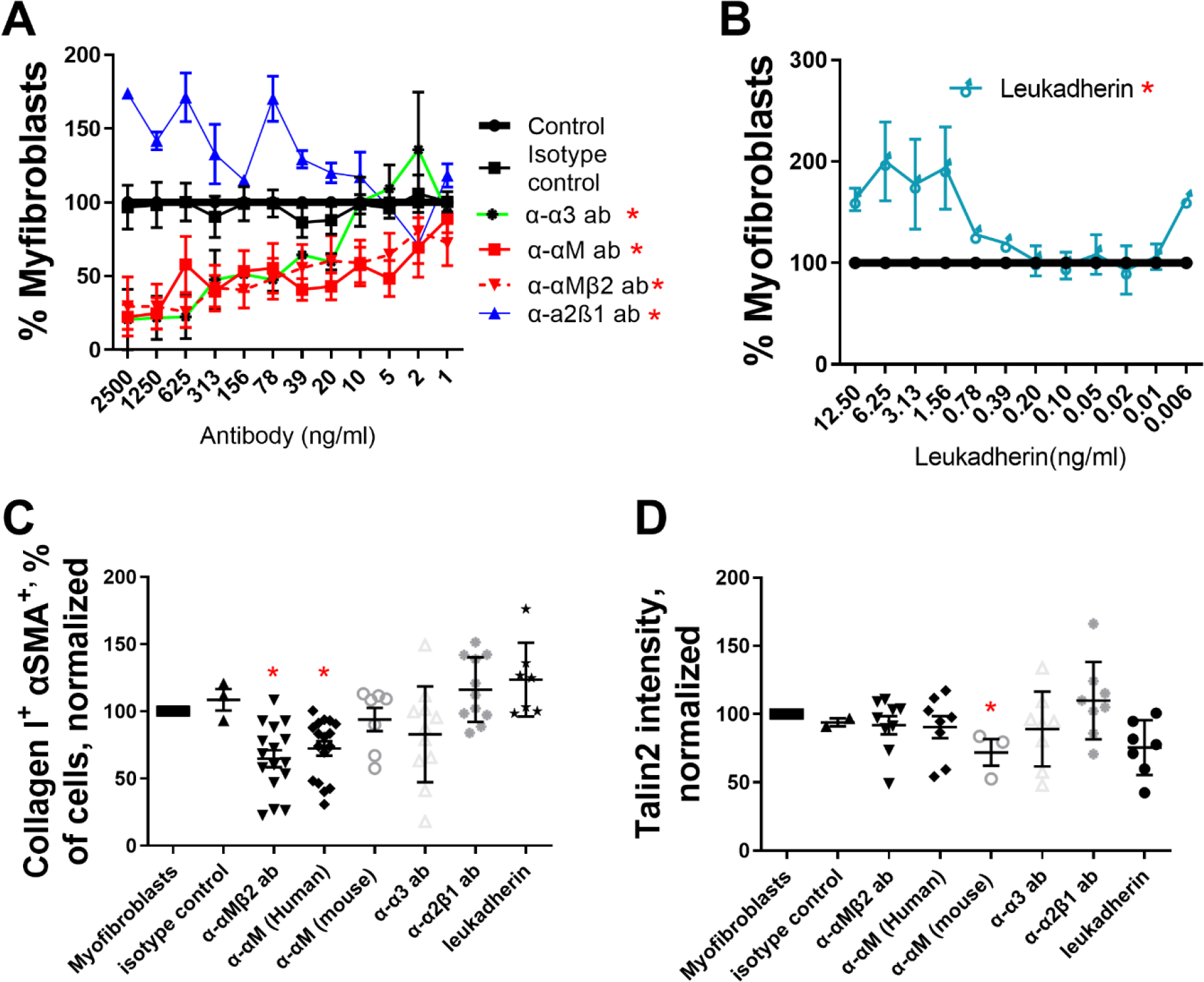
Antibodies against integrins α3, αM, and αMβ2 de-differentiate human myofibroblasts from monocyte precursors. Freshly isolated human monocytes (A-D) were differentiated into myofibroblasts and subsequently treated with (A) α-αMβ2, α-αM, α-α3, or α-α2β1 antibodies or (B) leukadherin at the indicated concentrations over 1 week, and the number of remaining myofibroblasts was assessed by morphology. Monocyte-derived myofibroblasts were treated with antibodies at 500 ng/ml and leukadherin at 2 ng/ml and analyzed via flow cytometry for (C) αSMA+ collagen I+ double positive cells and (D) talin2+. n ranges from 3 to 15. * = statistical significance of P < 0.05, < 0.01, or < 0.001, significance vs isotype control antibody, (A and B) 2-way ANOVA (with Fisher’s LSD post test), (C and D)

**Table 2:**
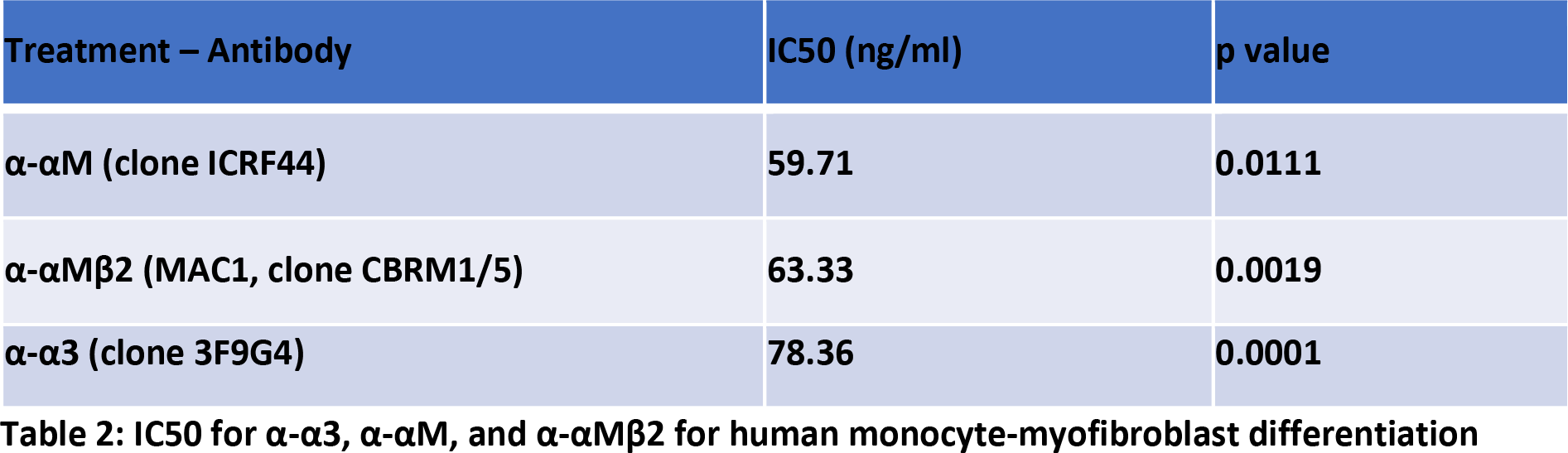
IC50 for α-α3, α-αM, and α-αMβ2 for human monocyte-myofibroblast differentiation

| Treatment – Antibody | IC50 (ng/ml) | p value |
| --- | --- | --- |
| α-αM (clone ICRF44) | 59.71 | 0.0111 |
| α-αMβ2 (MAC1, clone CBRM1/5) | 63.33 | 0.0019 |
| α-α3 (clone 3F9G4) | 78.36 | 0.0001 |

Taken together, these results show that our mRNAseq study revealed key integrins which can govern myofibroblasts differentiation and de-differentiation, when their ligand binding is promoted or inhibited.

To determine if we could reduce intracellular markers of myofibroblast differentiation, we added α-α3, α-αM, and α-αMβ2 to monocyte-derived myofibroblasts and measured the number of αSMA^+^ collagen I^+^ double positive cells via flow cytometry. The IC50 from Figures 1A and 1B was used to generate an effective dose of 500 ng/ml antibody and 2 ng/ml leukadherin. 500 ng/ml α-αM and α-αMβ2 reduced αSMA^+^ collagen I^+^ cells from myofibroblast precursors, indicating that de-differentiation of monocyte- myofibroblasts was not limited to morphological changes (Figure 1C). Similarly, both leukadherin and the integrin α2β1 stabilizing antibody increased the amount of αSMA^+^ collagen I^+^ double positive cells (Figures 1B and 1C).

In a companion study, we showed that adhesion to a surface of sufficient stiffness (>12 kPa) is essential for monocyte-myofibroblast differentiation, and that inhibition of the stiffness-sensing spring protein talin2 de-differentiated monocyte-derived myofibroblasts and reversed existing lung fibrosis in a mouse model [44]. Neither integrin blockers (α-α3, α-αM, and α-αMβ2) nor integrin interaction promoters (Gi14, leukadherin) significantly altered talin2 expression in monocyte-derived myofibroblasts (Figure 1D). Thus de-differentiation of monocyte-derived myofibroblasts by blocking antibodies (α-αMβ2, α-αM, α-α3) does not appear to operate through inhibition of talin2.

Myofibroblasts contribute to the development and maintenance of scar tissue in several distinct ways: by directly secreting ECM components (including collagen I), by using spindle-shaped morphology and cytoskeletal structures (FAs and FBs) to increase tissue rigidity, and by secreting pro-fibrotic cytokines and chemokines.

To determine if de-differentiation is accompanied by a loss of intracellular structures found in myofibroblasts (FAs and FBs), we added 500 ng/ml α-α3, α-αM or α-αMβ2 to cultured monocyte-derived myofibroblasts. α-α3, α-αM or α-αMβ2 eliminated myofibroblast’s spindle-shaped morphology and caused myofibroblasts to adopt a more macrophage-like morphology (Figure S1A vs B-D). α-α3, α- αM and α-αMβ2 also induced the complete loss of myofibroblast cytoskeletal structures (FAs and FBs, Figure S1B-D).

To determine if myofibroblast de-differentiation is accompanied by a change in secretome, we added 500 ng/ml α-α3, α-αM or α-αMβ2 to monocyte-derived-myofibroblasts. α-αM and α-αMβ2 reduced the amount of secreted pro-fibrotic macrophage-chemotactic protein-1 (MCP1) from monocyte-derived myofibroblasts (Figure S2A). α-αMβ2 also inhibited the amount of secreted IL-6 (Figure S2C). Antibody clone Gi14 (which stabilizes the association of integrin α2β1) and leukadherin (which stabilizes integrin αMβ2) both promoted the secretion of MCP1 (Figure S2A) and IL-6 (Figure S2C) in monocyte-derived myofibroblasts, again showing that modulation of specific integrins can both promote or de- differentiate myofibroblast morphological phenotype and secretome of myofibroblasts.

To determine if modulation of integrins could de-differentiate mouse myofibroblasts, we added α-α3, α- αM, α-αMβ2, α-α2β1 (Gi14) and leukadherin to mouse monocyte-service myofibroblasts and mouse fibroblast-derived myofibroblasts. Blocking antibodies cross-reactive against mouse integrins α3, αM, αMβ2 (α-α3, α-αM, α-αMβ2) de-differentiated mouse monocyte-derived myofibroblasts (Figure 2A).

**Figure 2:**
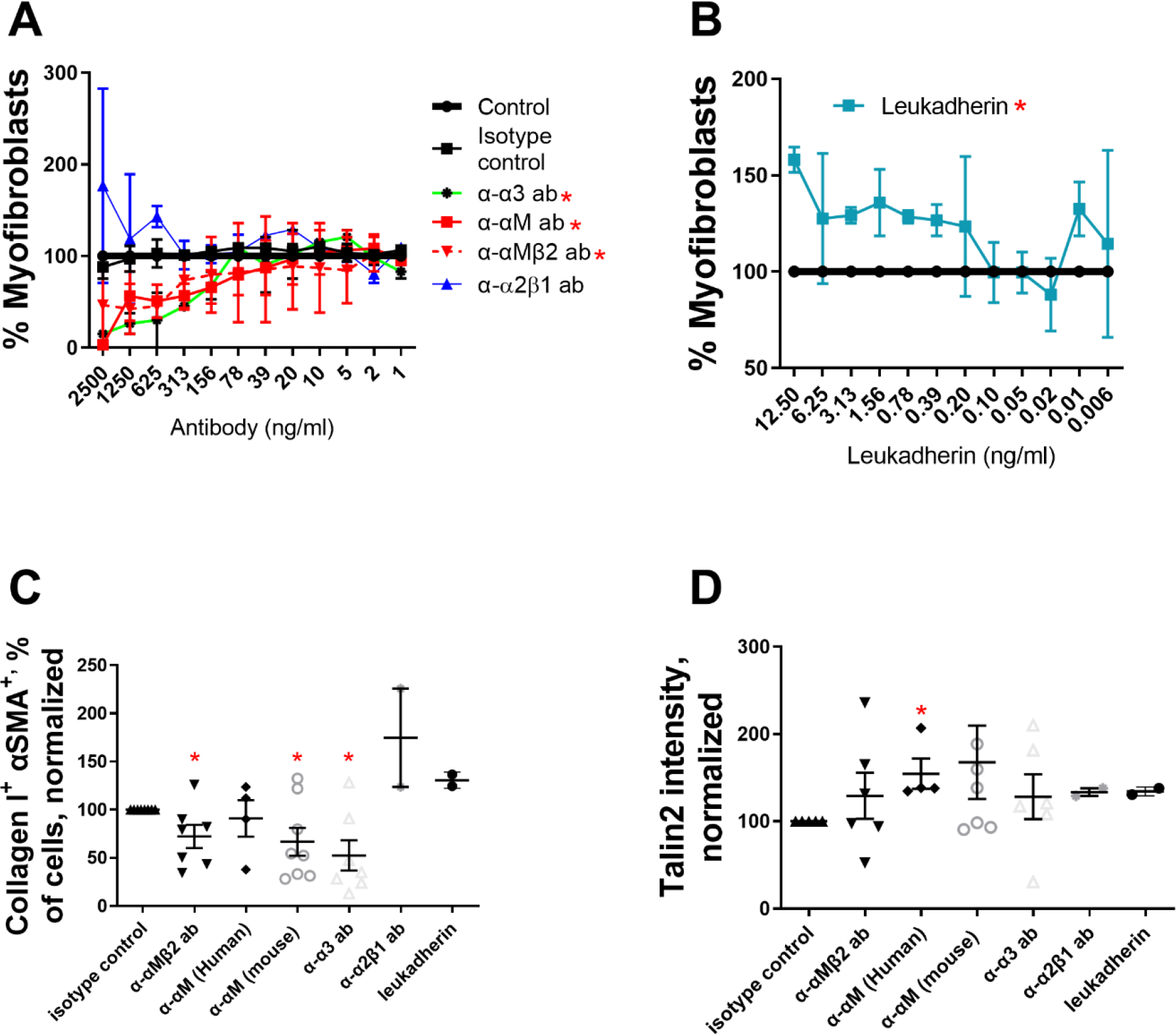
Antibodies against integrins α3, αM, and αMβ2 de-differentiate mouse myofibroblasts from monocyte precursors. Freshly isolated mouse monocytes (A-D) were induced to become myofibroblasts with IL-13, MCSF, and β-mercaptoethanol, treated with (A) α- αMβ2, α-αM, α-α3, or α-α2β1 antibodies or (B) leukadherin at the indicated concentrations over 1 week, and the number of myofibroblasts was assessed by morphology. Monocyte-derived myofibroblasts were treated with antibodies at 500 ng/ml and leukadherin at 2 ng/ml, allowed to de-differentiate, and analyzed via flow cytometry for (C) αSMA+ and collagen I+ double positive cells and (D) talin2+. n ranges from 2 to 7. * = statistical significance of P < 0.05, < 0.01, or < 0.001, significance vs isotype control antibody, (A and B) 2-way ANOVA (with Fisher’s LSD post test), (C and D) Student’s t-test.

Promoting the association of integrin α2β1 (via Gi14) and αMβ2 (via leukadherin) both increased monocyte-derived myofibroblast differentiation (Figure 2A and B). α-α3, α-αM, α-αMβ2 also reduced the number of αSMA^+^ and collagen I^+^ double positive myofibroblasts (Figure 2C), and did not decrease the amount of talin2^+^ cells (Figure 2D). This again suggests that de-differentiating myofibroblasts by blocking α3, αM, αMβ2 operates through a different mechanism than de-differentiating myofibroblasts by inhibition of talin2. Stabilizing integrin α2β1 (through Gi14) and αMβ2 (via leukadherin) increased both the amount of α αSMA^+^ and collagen I^+^ double positive monocyte-myofibroblasts, as well as increasing talin2 concentration in these cells (Figures 2C and 2D).

Antibody CBRM1/5 is raised against the activation epitope of human αMβ2 [47], but shares 2 amino acid (AA) overlap with mouse αMβ2 [48]. However, CBRM1/5 de-differentiates mouse monocyte-derived myofibroblasts (Fig 2A), though at a reduced effectiveness vs human monocyte-derived myofibroblasts (Fig 1A). While it would have been ideal to use an anti-mouse αMβ2 antibody, no such antibody exists raised against the active conformation of mouse αMβ2.

To determine if de-differentiation of mouse myofibroblasts is accompanied by an altered secretome, we assessed the conditioned media from mouse myofibroblasts treated with 500 ng/ml α-α3, α-αM, and α-αMβ2. De-differentiation of myofibroblasts did not alter the amount of secreted anti-fibrotic IL-10, but in some cases significantly reduced (and did not increase in any case) the amount of pro-fibrotic IL-23 [49], CCL22 [50], IL-6 [51], CCL17 [50, 52], IL-12 subunit p40 [53], CXCL1 [54], and TNF-α [55] (Figure S3).

While monocytes can become myofibroblasts, the primary cellular component of scar tissue in fibrosis is the fibroblast-derived myofibroblast. To determine if fibroblast-derived myofibroblasts could be de- differentiated, we added 500 ng/ml α-αMβ2, α-αM, α-α3 to fibroblast-derived myofibroblasts and measured the number of αSMA^+^ and collagen I^+^ double positive cells, and talin2^+^. α-α3 reduced the number of αSMA^+^ and collagen I^+^ fibroblast-derived myofibroblasts, while α-αM and α-αMβ2 did not (Figure S4A), consistent with the expression of these integrins on monocytes but not fibroblasts. Neither α-αMβ2, α-αM, α-α3 treatment lowered the amount of talin2 (Figure S4B), again confirming that de- differentiation of myofibroblasts by inhibiting integrin binding and inhibiting tension sensing operate by non-overlapping mechanisms. α-α3 decreased the amount of IL-6 secreted from fibroblast-derived myofibroblasts (Figure S2D). No antibody treatment lowered the number of mouse αSMA^+^ and collagen I^+^ double-positive fibroblast-derived myofibroblasts, or talin2^+^ (Figures S4C and D).

Treatment with antibodies did not induce cell death from monocyte (Figure S5A and C) or fibroblast populations (Figure S5B and D), among both human and mouse cells. This confirms that the de- differentiation of myofibroblasts is not caused by, or accompanied by, an increase in cell death.

Conjugation of decorin’s collagen-binding peptide (CBP) to an antibody increased the proportion of the conjugated CBP-antibody that reached and was retained in fibrotic lungs, compared to non-conjugated antibody [43]. This finding involved antibodies against soluble factors (α-TNFα and α-TGFβ). To determine if we could deliver an anti-integrin antibody to a fibrotic organ, we conjugated CY7-CBP to α- αM. We instilled mouse lungs with bleomycin, allowed fibrosis to develop over a week, and injected Cy7-CBP-α-αM and Cy7-α-αM i.v. in mice with healthy and fibrotic lungs. We compared the fluorescence of the harvested organs after 48 hr via IVIS (Figure 3A). Direct comparison of the fluorescence of healthy and fibrotic lungs (Figure 3B) shows that significantly more fluorescently labeled antibody remained in fibrotic lungs after 48 hr. Because αM is present on monocytes, and monocytes are enriched in the spleen, we compared the fluorescence of spleens (Figure 3C). There was significantly less Cy7-CBP-α-αM in the spleen in animals with fibrotic lungs, suggesting that the enrichment of Cy7-CBP-α-αM in the lungs was coming partially at the expense of Cy7-CBP-α-αM in the spleen.

**Figure 3:**
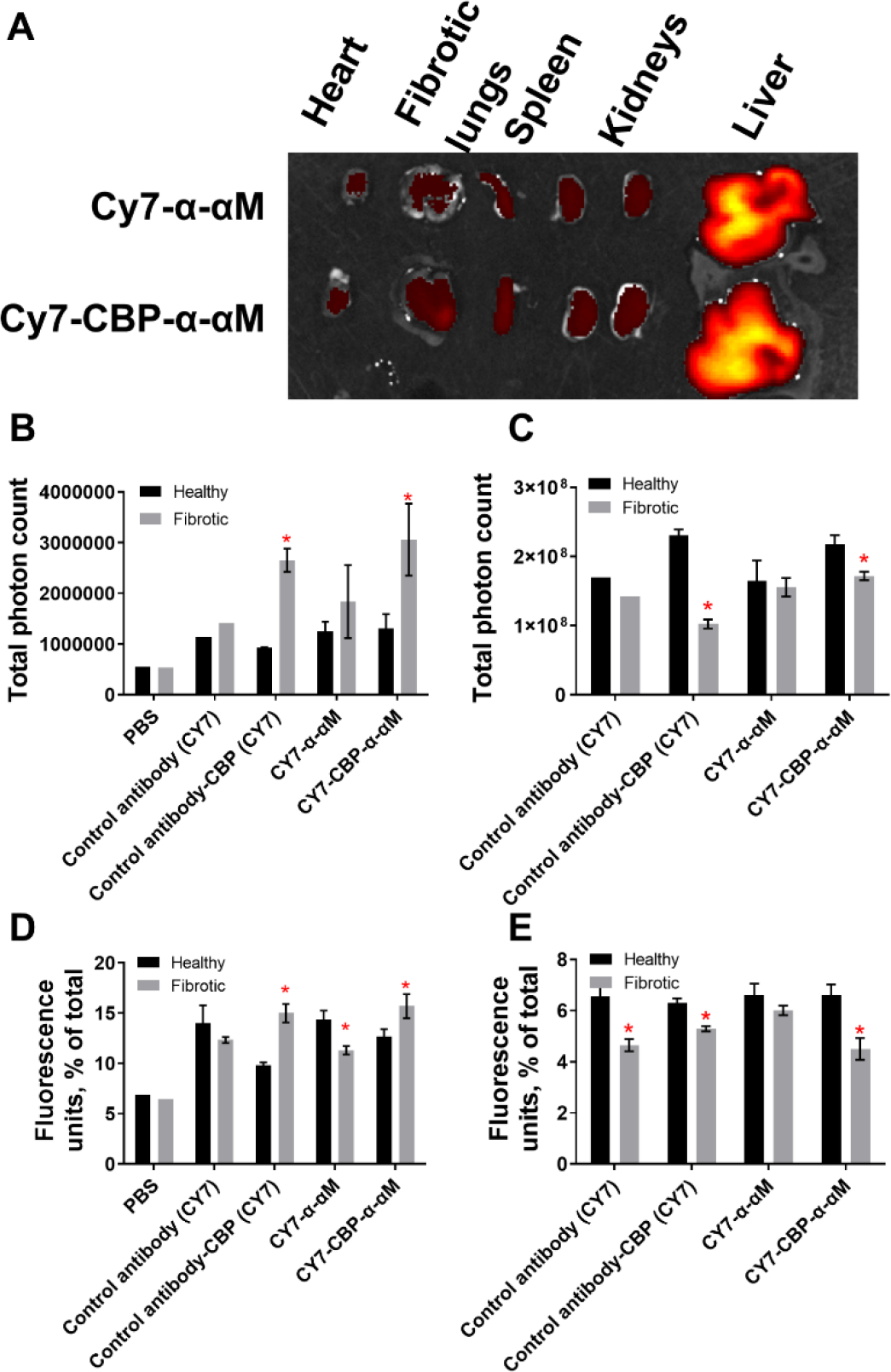
Conjugation of decorin’s collagen-binding peptide (CBP) to anti-integrin- αM (α-αM) increases antibody concentration in fibrotic lungs. Mice were intranasally instilled with 75 μg of bleomycin sulfate (fibrotic) or PBS (healthy) and injected 1 week later with Cy7- α-αM or Cy7-CBP-α-αM. (A) Heart, lung, spleen, kidneys, and liver were harvested 48 hr after injection, and fluorescence intensity was measured via IVIS. (B) The number of photons from lungs or (C) spleen. Fluorescence of the resected organs was pooled and the percentage of fluorescence associated with (D) lungs and (E) spleen. n ranges from 2 to 4. * = statistical significance of P < 0.05, < 0.01, or < 0.001, significance is fluorescence of fibrotic lungs vs fluorescence of healthy lungs, Student’s t-test.

To make certain that our Cy7-CBP-α-αM is not simply remaining in circulation in the mouse for longer, we pooled the total fluorescence of all organs for Cy7-CBP-α-αM and CY7-CBP conjugated isotype control antibody, and normalizing the fluorescence for each organ. This analysis showed significantly more Cy7-CBP-α-αM in fibrotic lungs than Cy7-α-αM in fibrotic lungs or Cy7-CBP-α-αM in healthy lungs (Figure 3D) and less in the spleen (Figure 3E).

To determine if we could reverse existing fibrosis in addition to de-differentiating myofibroblasts, we conjugated non-fluorescent CBP to each of α-α3 (CBP-α-α3), α-αM (CBP-α-αM), and α-αMβ2 (CBP-α- αMβ2). Using our recently published results as a guide [43], we used different molar excess of CBP to establish ratios sufficient to attach at least 5 CBP peptides (on average) to each antibody (Figure S6).

We instilled mouse lungs with bleomycin, allowed fibrosis to develop, and injected α-α3, α-αM, α-αMβ2, CBP-α-α3, CBP-α-αM, and CBP-α-αMβ2 at 7, 9, 11, 14, 16, and 18 days post bleomycin insult. The mice were euthanized 21 days post insult. Only CBP-α-α3 and α-αMβ2 provided weight gain that was statistically higher than the no-treatment control (Figure 4A-C). CBP-α-α3, α-αM, CBP-α-αM, α-αMβ2, and CBP-α-αMβ2 significantly reduced the total amount of collagen in the right lobe of the lung vs the untreated fibrotic lungs, as assessed by hydroxyproline assay (Figure 4D). CBP-α-αM and CBP-α-αMβ2 significantly reduced the amount of collagen in the right lungs as a percentage of overall lung weight, compared to the untreated fibrotic lungs (Figure 4E).

**Figure 4:**
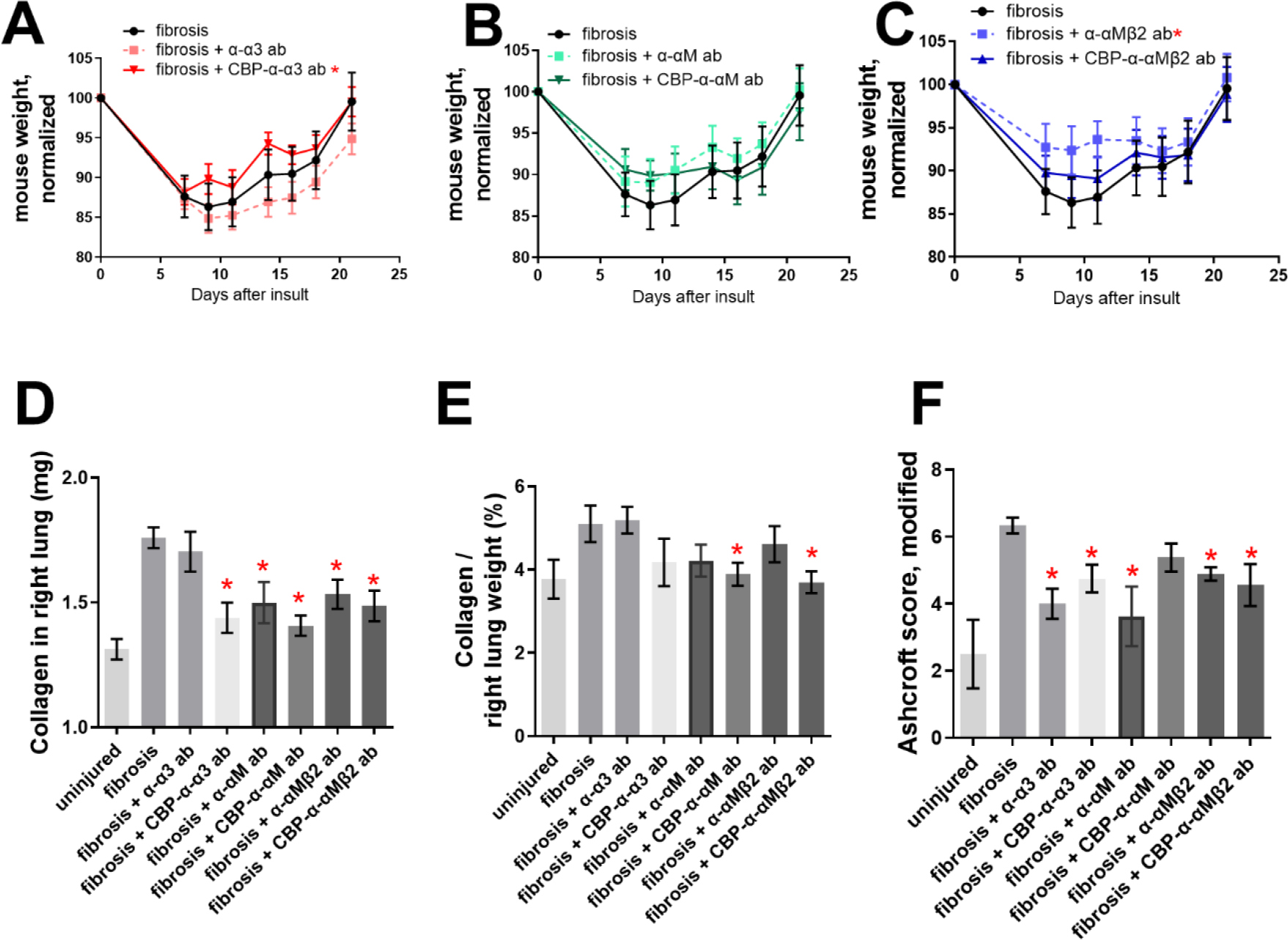
Antibodies against integrins α3, αM, and αMβ2 rescues the fibrotic damage from bleomycin insult to mouse lungs. 50 µg antibody was injected i.v. after 7, 9, 11, 14, 16, and 18 days following insult by 75 μg bleomycin. (A-C) mouse weights after treatment with (A) α-α3 and CBP-α-α3, (B) α-αM and CBP-α-αM, (C) α-αMβ2 and CBP-α-αMβ2. (D) Collagen content from the right, multi-lobed lung assessed by hydroxyproline assay. (E) Data from D divided by dry weight of right lobes of mouse lungs. (F) Blinded Ashcroft scoring. n ranges from 8 to 9. * = statistical significance of P < 0.05, < 0.01, or < 0.001, significance vs fibrotic lungs (A-C) 2-way ANOVA (with Fisher’s LSD post-test), (D-F) Student’s t-test.

The left lobes of the bleomycin-insulted lungs were mounted in paraffin, sectioned, stained using Masson’s trichrome, and the histology was blindly scored using the modified Ashcroft method [56] by a researcher not involved in administration of the treatment (Figure 4F). Both α-α3 and CBP-α-α3 reduced the Ashcroft score, as did both α-αMβ2 and CBP-α-αMβ2; with the α-αMβ2 antibody, only the unmodified antibody showed a reduction (Figure 4F-G). Representative images of the Masson’s trichrome stained left lobes can be seen in Figure 5 A-H.

**Figure 5:**
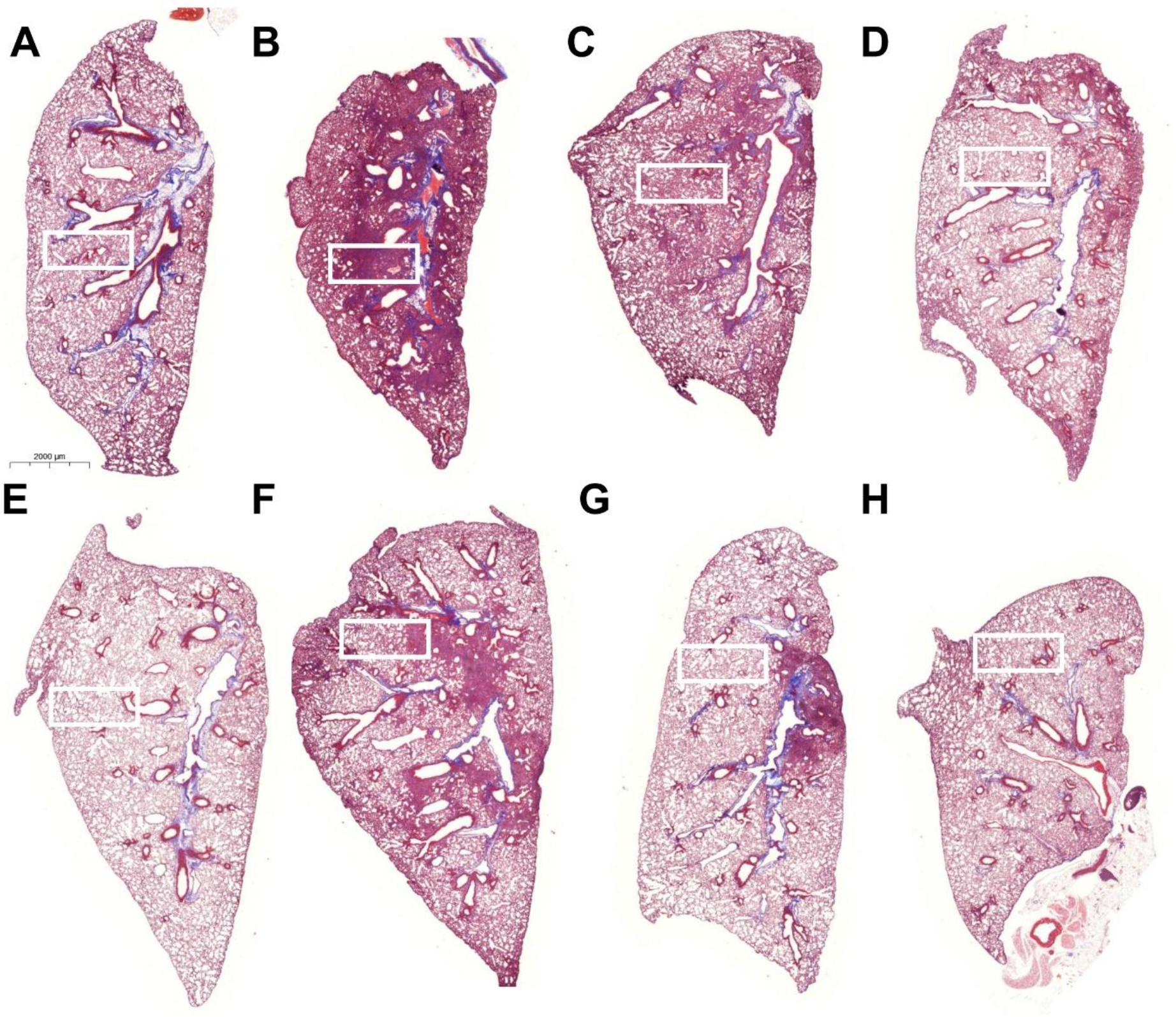
Representative images of left, single-lobed lungs stained with Massons’s trichrome. (A) Uninsulted lungs instilled with PBS. Lungs insulted with (B-H) 75 μg bleomycin, and instilled with (C) α-α3, (D) CBP-α-α3, (E) α-αM, (F) CBP-α-αM, (G) α-αMβ2, (F) CBP-α-αMβ2. Lungs were harvested 3 weeks after insult and stained via Masson’s trichrome. Scores in Figures 4, 6, and inset images in supplemental figure 7.

To determine if treatment by α-α3, α-αM, and α-αMβ2 and CBP-α-α3, CBP-α-αM, and CBP-α-αMβ2 improved fibrosis in both male and female mice, we analyzed the hydroxyproline-based quantitative collagen data (Fig 6A-D) and histology-based Ashcroft qualitative data (Fig 6E-H) based on mouse sex. In general, in this model males show a more fibrotic response. Treatment by α-α3, α-αM, and α-αMβ2 and CBP-α-α3, CBP-α-αM, and CBP-α-αMβ2 reduced the absolute amount of collagen in the right lobes of both male and female mice (Figure 6A-B), though reductions in the amount of collagen as a portion of lung weight were not significant except in the case of CBP-α-αMβ2 in female mice (Figure 6C-D). Several treatments improved the modified Ashcroft scores for both male and female mice (Figure 6EH).

**Figure 6:**
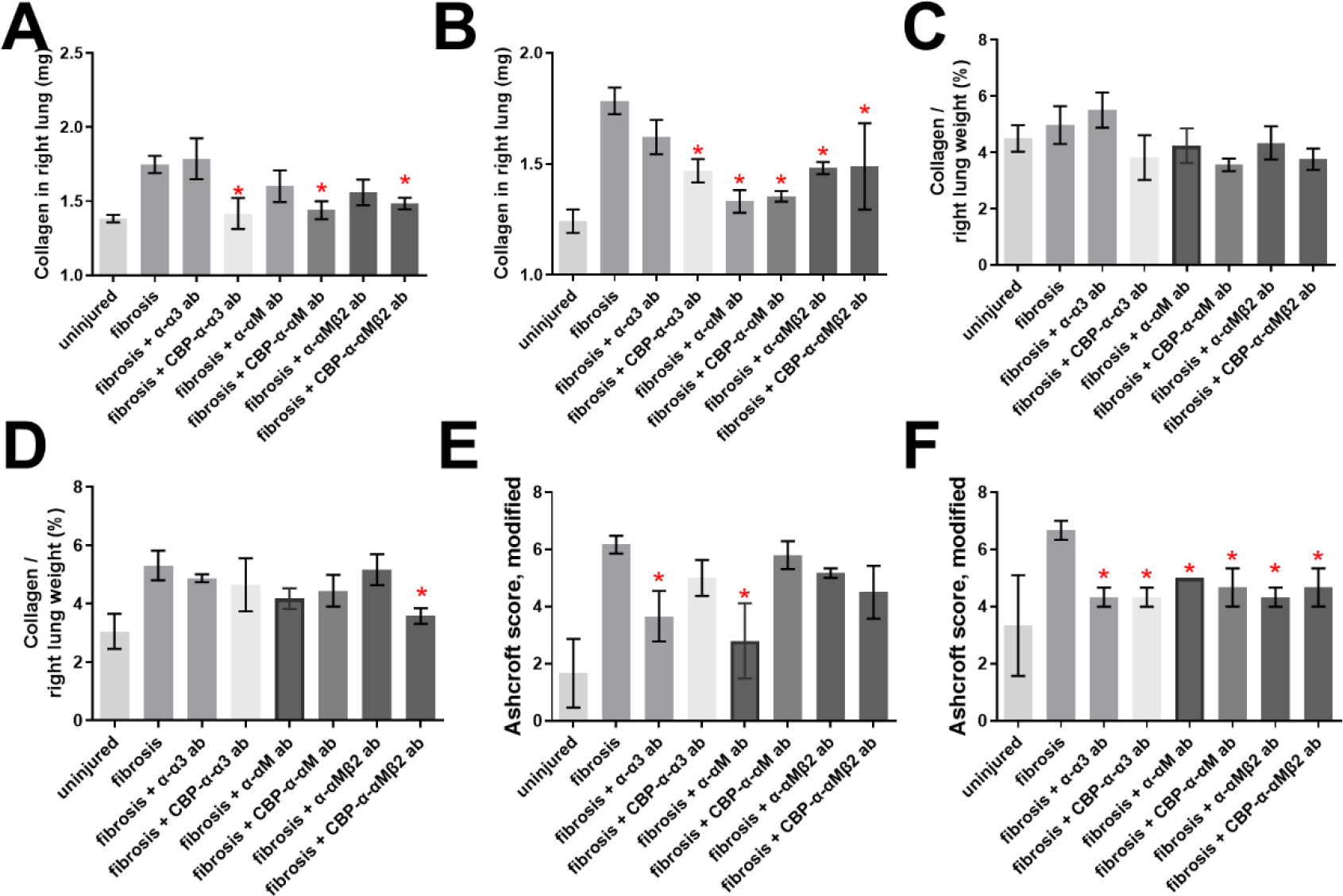
Antibodies against integrins α3, αM, and αMβ2 rescue the fibrotic damage from both male and female mice. Data from Figure 4, treatment by α-α3, CBP-α-α3, α-αM, CBP-α-αM, α- αMβ2, CBP-α-αMβ2 compared across male (n=5 or 6, A, C, E, G) and female (n=3, B, D, F, H) mice for amount of collagen in lungs (A, B), Collagen as a percentage of dry lung weight (C, D), Ashcroft scores (E, F), and blinded Ashcroft scores (E, F). n ranges from 3 to 6. * = statistical significance of P < 0.05, < 0.01, or < 0.001, significance vs fibrotic lungs unless otherwise indicated, Student’s t-test.

Overall, these data indicate that treatment of fibrosis with blocking anti-integrin antibodies directed against α3 and αMβ2 can reverse pulmonary fibrosis after a bleomycin insult.

To determine if we could reverse existing kidney fibrosis in addition to existing lung fibrosis, we attempted to target CBP-tagged antibodies to fibrotic kidneys, similar to Figure 3 for lungs. We ligated the descending ureter of the left kidney (unilateral ureteral obstruction, UUO) and allowed the kidneys to become damaged for 1 week. We then injected Cy7-CBP-α-αM and Cy7-α-αM i.v. We compared the fluorescence of the harvested organs after 48 hr via IVIS (Figure 7A). As in Figure 3, to make certain that Cy7-CBP-α-αM is not simply remaining in circulation in the mouse for longer, we pooled the total fluorescence of all organs for Cy7-CBP-α-αM and CY7-α-αM, and normalizing the fluorescence for each organ. This analysis showed significantly more Cy7-CBP-α-αM in fibrotic kidneys than in healthy kidneys, in comparison to the ratio for Cy7-α-αM (Figure 7B). We also tested the targeting potential of Cy7-CBP- α-TGFβ, which we had previously shown to be anti-fibrotic when targeted to lungs (Figure 7B) [43].

**Figure 7:**
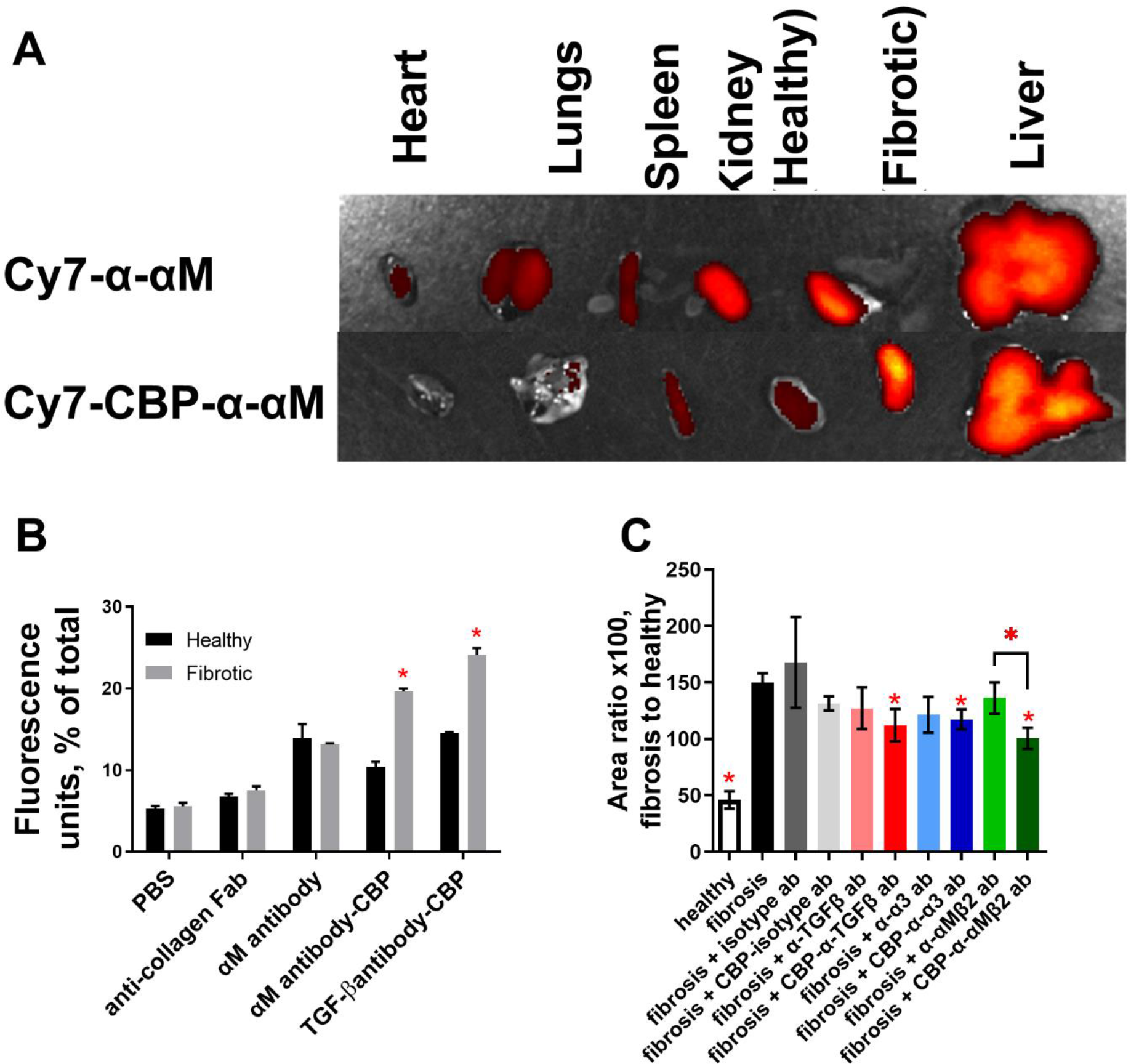
Antibodies against TGFβ, integrin α3, and integrin αMβ2 rescue the fibrotic damage from UUO insult to mouse kidneys. The descending ureter of the left kidney was surgically ligated, and 50 µg antibody (either tagged with cy7 or unconjugated) was injected i.v. after 7 days. (A) Heart, lung, spleen, kidneys, and liver were harvested 24 hr after injection, and fluorescence intensity was measured via IVIS. (B) Fluorescence of the resected organs was pooled and the percentage of fluorescence associated with healthy and fibrotic kidneys. (C) kidneys were resected 14 post UUO insult (7 days post antibody injection), mounted, and stained via immunohistochemistry for collagen I. The amount of positive IHC staining was compared to the overall amount of kidney tissue, per image. For A and B, n=3. For C, healthy n =10, fibrosis n= 10, CBP-antibody treated kidneys n = 8, and unconjugated-antibody treated kidneys n=6. * = statistical significance of P < 0.05, < 0.01, or < 0.001, (B) significance is fluorescence of fibrotic left kidney vs fluorescence of healthy kidney, Student’s t-test. (C) ANOVA vs fibrosis control, Welch’s correction. Comparison between α-αMβ2 and CBP-α-αMβ2 is Student’s t-test.

These results indicate we can target our antibodies to fibrotic kidneys, and in greater quantity than to fibrotic lungs.

To determine if CBP-α-TGFβ, CBP-α-α3, and CBP-α-αMβ2 could reverse existing kidney fibrosis, we performed a UUO on the left kidney. At 1 week post-UUO we injected 50 micrograms of α-TGFβ, α-α3, α-αMβ2, CBP-α-TGFβ, CBP-α-α3, and CBP-α-αMβ2 i.v. At 2 weeks post-UUO, we resected the kidneys, and assessed fibrosis via immunohistochemistry (IHC) for collagen I (Figure 7, Figure S8) and blood markers of kidney injury (Figure S9). CBP-α-TGFβ, CBP-α-α3, and CBP-α-αMβ2 lowered the amount of collagen present in the UUO kidneys (Figure S8, Figure 7C). An analysis of blood taken from the kidney UUO mice indicates that each antibody treatment reduced the amount of circulating creatine kinase, but did not change the concentration of blood urea nitrogen (BUN) or uric acid (Figure S9). This could be because the UUO model leaves mice with one entirely healthy kidney. These results indicate a single 50 microgram dose of CBP-α-TGFβ, CBP-α-α3, and CBP-α-αMβ2 is capable of partially reversing kidney fibrosis.

In summary, α-α3, α-αM, and α-αMβ2 were identified as proteins upregulated during the differentiation of monocytes into myofibroblasts. Blocking antibodies against α-α3, α-αM, and α-αMβ2 reverse myofibroblast differentiation and reduce the pro-fibrotic secretome of myofibroblasts. Injected CBP- conjugated antibodies preferentially localize at fibroses in both the lungs and the kidneys, and CBP- conjugated blocking antibodies (CBP-α-TGFβ, CBP-α-α3, CBP-α-αM, and CBP-α-αMβ2) reverse lung and kidney fibroses at lower doses than untargeted antibodies. Transcripts for integrins α3, αM, and β2 are each upregulated in macrophages in idiopathic pulmonary fibrosis (IPF) (Figures S10, S11, and S12, data from IPF cell atlas [57]). These results raise the possibility of a targeted immunotherapy treatment for fibrosis.

## Discussion

Our overall objective in this work is to explore an approach by which to de-differentiate myofibroblasts and reverse fibrosis. This study was guided by protein targets identified from an RNAseq comparison of myofibroblasts under anti- and pro- fibrotic culture conditions related to surface stiffness (1 kPa and 12 kPa) [44]. These targets were subsequently validated and refined by *in vitro* experimentation, where only blocking antibodies α-αMβ2, α-αM, α-α3 consistently de-differentiated myofibroblasts (Fig 1 and 2, S1-S3). Each of α-αMβ2, α-αM, and α-α3 reversed existing fibrosis in mouse models of lung and kidney fibrosis (Fig 4-7).

Myofibroblast de-activation has long been a goal of fibrosis research. De-activation can be achieved by myofibroblast apoptosis [14] or de-differentiation [9]. Since no apoptosis was observed by immunofluorescence (Figure S1B-D) and no increase in live-dead staining was observed after treatment with α-α3, α-αM, or α-αMβ2 (Fig S5), we conclude we have de-activated myofibroblasts through de- differentiation. De-activating monocyte-derived myofibroblasts by de-differentiation seems preferable to apoptosis, since monocyte-derived cells are proficient at removing deposited ECM and regenerating tissue, both of which are required to reverse fibrosis [7].

While we showed in this paper that targeting the αMβ2 complex and αM both de-differentiate myofibroblasts and reverse fibrosis, targeting β2 (CD18) with blocking antibodies resulted in apoptosis of myofibroblasts [45] (Data not shown), which rendered this blocking antibody unsuitable for use in this study. However, this reported result may be an example of de-activation of myofibroblasts by apoptosis.

While we have shown that α-αM and α-αMβ2 reverse existing fibrosis, some evidence indicates they may have a protective role against fibrosis as well. α-αM is effective in reducing tissue damage in a kidney-ischemia model [58], while α-αMβ2 antibody prevented both damage from kidney-ischemia and subsequent ischemia-induced fibrosis [59].

Only α-α3—and not α-αM or α-αMβ2—de-differentiated fibroblast-derived myofibroblasts (Figure S4). While fibroblasts express integrin α3, the integrins αM and αMβ2 are considered canonical myeloid markers. Thus, it appears that anti-integrin antibodies are sufficiently specific that different lineages of myofibroblasts can be targeted by anti-integrin antibodies to reverse fibrosis.

Hinz has proposed the “super-mature” FAs are essential for the development and maintenance of the myofibroblast phenotype [9]. Super-mature FAs concentrate in one location several different protein components of cell adhesion and tension sensing: integrins, tension-sensing talins, and cytoskeletal machinery [31, 32]. FAs participate in inside-out and outside-in signaling in relation to their size [60]. Figure S1A (inset) shows structures remarkably similar to Hinz’s super-mature FAs [29], and images in Figure S1B, C and D show that treatment of myofibroblasts with α-αMβ2, α-αM, and α-α3 both remove both the myofibroblast morphology and the presence of FAs and FBs. That removal of the myofibroblast morphology occurs in parallel with the removal of FAs and FBs perhaps confirms Hinz’s proposal that super-mature FAs are critical structures for the maintenance of myofibroblasts [9].

αMβ2 is necessary for maintenance of monocyte binding, allowing the actin reorganization that sustains adhesion [61]. This suggests that disruption of αMβ2 might remove existing FAs, again recalling Hinz’s hypothesis that FAs are critical to myofibroblast differentiation and maintenance. Treatment of monocyte-derived myofibroblasts with α-αMβ2, α-αM and α-α3 induced a morphology (Figure S1B-D) that appears similar to fluorescence images from monocytes [62] and fibroblasts [63, 64] that have had talin2 reduced, suggesting that integrin- blocking antibodies and inhibition of talin2 both might de- differentiate myofibroblasts through disruption of existing FAs.

Talins and integrins interact in multiple ways within FAs. Talin’s interaction with the β-tail of integrins [24] is essential for “outside-in” signaling of integrins [65]. Talin2’s affinity for the β-tail of β2 increases the binding of monocytes to cells in two key examples. First, talin2’s interaction with β2 promotes αLβ2 adhesion to ICAM-1 [66]. Second, talin2 is also essential for αMβ2-mediated phagocytosis [62]. The differences in tension-sensing and mechanotransduction between talin1 and talin2 may be primarily due to each isoform’s respective integrin associations [25], suggesting the interaction of talins and integrins may play a large role in tension sensing. Intriguingly, integrin-dependent mechanosensing is talin isoform specific [64]. The difference in talin’s affinity for each integrin [25], the subcellular localization of integrins and talins, and interactions with the adhesome are all part of the dynamic mechanosensing and mechanotransduction of monocytes during myofibroblast differentiation [8].

While integrin interactions with tension-sensing talins appears to be key to the formation and maintenance of myofibroblasts, integrins themselves may play a role in tension sensing. The speed at which integrin-actin linkages are formed and broken has been suggested to be a cellular tension sensing mechanism in and of itself [33, 67–69]. This suggests that integrins may transduce information about specific binding ligands (including ECM components) and surface stiffness. This may explain how several integrins appeared in a screen designed to assess the differences in cultured monocytes between soft (1 kPa) and stiffer (12 kPa) surfaces, on the stiffer surfaces permitting monocyte-to-myofibroblast differentiation.

De-differentiation of mouse and human myofibroblasts resulted in strongly reduced IL-6 and MCP-1 secretion (Figure S2 and S3). Removing the MCP-1 pathway abrogates kidney injury [70], again suggesting monocyte-derived cells are quite important in fibrosis. IL-6 is quite pro-fibrotic, in some cases sufficient to induce myofibrobasts [71]. That MCP-1 and IL-6 were the two consistently inhibited secreted proteins in the cytokine screens suggests that de-differentiation of monocyte-derived myofibroblasts may have additional effects beyond the de-activation of individual myofibroblasts, and may contribute to an overall reduction in the pro-fibrotic environment in general.

Several anti-integrin therapies have failed to translate to medicinal use due to side-effects or lack of efficacy [24]. Our hope was that by targeting antibodies to fibrotic tissue using the decorin-derived CBP (Fig 3 and 7), we could increase the local concentration of anti-integrin antibodies in fibrotic organs, and potentially make anti-integrin antibodies a more useful therapeutic. We further tested a monoclonal antibody that recognizes only the activation epitope of αMβ2 [47], clone CBRM1/5. This antibody binds only to a conformational epitope exposed on activated αMβ2. By using an antibody that recognizes only a conformational epitope, we hope to further reduce the side-effects of anti-integrin treatments.

Each of α-α3, α-αM, and α-αMβ2 (and their CBP-conjugated variants) de-differentiated mouse myofibroblasts and reversed fibrosis in mice at an identical dose to previously published antibody therapeutics for fibrosis [43]. This indicates that the local concentration of untargeted α-α3, α-αM, and α-αMβ2 is sufficient to reverse fibrosis in mice. CBP-conjugated antibodies were twice as concentrated as unconjugated antibodies in fibrotic lungs after 48 hr (Fig 3B) and in fibrotic kidneys after 24 hours (Fig 7B). A single 50 microgram dose of CBP-conjugated antibodies was sufficient to partially reverse fibrosis in a kidney model of unreversed UUO kidney fibrosis (Figure 7 and S8).

Our results show that α-α3, α-αM, and α-αMβ2 may be potentially useful as translational therapeutics, capable of bringing immunotherapy treatments to fibrosing diseases. We again note that the α-αMβ2 clone used was less effective on mouse myofibroblasts (and thus presumably in the mouse model) than on human monocyte-derived myofibroblasts. Clinical attractiveness may be particularly high for CBP-α-αMβ2 (clone CBRM1/5), which both recognizes only the active ligand-binding configuration for αMβ2, and is also capable of being targeted to sites of fibrosis to increase the local concentration.

## Materials and Methods

### Study design

This study was designed to test the strategy that key pathways of myofibroblast differentiation can be revealed by an RNAseq of myofibroblast precursors (monocytes) cultured on anti-fibrotic soft (1 kPa) and pro-fibrotic stiff (12 kPa) surfaces [44]. Specifically, this study examines whether antibodies against key integrins upregulated in monocyte-myofibroblasts (α-α3, α-αM, and α-αMβ2) can de-differentiate myofibroblasts and reverse fibrosis in a mouse model of lung fibrosis. CBP functionalization was employed to enhance retention in the fibrotic microenvironment.

Group size was selected based on experience with the pulmonary fibrosis model. Mice were randomized into treatment groups within a cage to eliminate cage effects from the experiment. Treatment was performed by multiple researchers over the course of this study, to ensure reproducibility. Lungs were also resected by multiple researchers, and blinded scoring was used on the fibrosis histology images.

### Purification of human monocytes

All human blood was acquired through the University of Chicago blood donation center, in accordance with human subject protocol at the University of Chicago. In order to isolate more peripheral blood mononuclear cells (PBMCs) than is possible through a single blood donation, PBMCs were purified from leukocyte reduction filters from de-identified blood donors. Blood was filtered, and leukocytes purified, the same afternoon as a morning blood donation, to reduce the amount of time that PBMCs could adhere to the filter.

Leukocyte reduction filters were sterilized with 70% ethanol, and the blood flow tubes on either end were clamped shut. The tube through which filtered blood had exited the filter was cut below the clamp, and a syringe containing 60 ml phosphate buffered saline (PBS) was inserted into the tube.

Following this, the tube through which unfiltered blood had entered the filter was unclamped, and PBS was slowly pushed through the filter in the opposite direction of the original blood flow. Reversing the flow of PBS through the filter resulted in a recovery of approximately 300 million cells per filter.

The collected blood was layered with lymphocyte separation media (LSM), and centrifuged at 1300 xG for 20 min. The PBMC layer was then removed by pipetting.

Monocytes were purified from PBMCs by use of a negative selection kit for human monocytes (Stemcell, Cambridge, MA), per the manufacturer’s instructions. Approximately 20 million monocytes were purified from each filter. Monocytes were then washed by PBS using five successive 300 xG centrifugation steps, in order to remove EDTA from the resulting population. Monocytes were checked for purity using flow cytometry, and average purity was above 95%. Monocytes were cultured immediately following purification, at 100,000 monocytes/cm^2^.

### Purification of mouse monocytes

All mouse experiments were performed under supervision with protocols approved by the University of Chicago IACUC. Spleens were resected from healthy C57BL/6 mice, pooled, and were placed in PBS with 1 mM EDTA, 2% fetal calf serum (FCS) to prevent subsequent clumping of cells. All PBS, plasticware, filters, glassware and magnets were pre-chilled to 4C, and kept cold throughout this procedure, to limit the clumping of cells. Importantly, ACK lysis buffer caused cell death, and so was not used in this procedure.

Spleens were pushed through a 100 μm filter to disassociate the cells. Monocytes were purified from disassociated cells by use of a negative selection kit (Stemcell), following the manufacturer’s instructions. The purified monocytes were then washed by PBS using five successive 300 xG centrifugation steps, in order to remove EDTA from the resulting population. Monocytes were checked for purity using flow cytometry, and average purity was above 95%. The average yield was 1.5 million monocytes per spleen.

Monocytes were cultured immediately following purification, at 250,000 monocytes/cm^2^.

### Culture of human and mouse monocytes

Human and mouse monocytes were cultured as previously described, using serum-free media (SFM) [72]. Briefly, SFM for human cells is composed of fibrolife (Lifeline, Frederick, MD), with 1x ITS-3 (Sigma, St. Louis, MO), 1x HEPES buffer (Sigma), 1x non-essential amino acids (Sigma), 1x sodium pyruvate (Sigma), and penicillin-streptomycin with glutamate (Sigma). For mouse monocytes, 2x concentrations of ITS-3, HEPES buffer, non-essential amino acids, and sodium pyruvate were added, with 50 μM β- mercaptoenthanol (ThermoFisher) and pro-fibrotic supplements M-CSF (25 ng/ml, Peprotech, Rocky Hill, NJ) and IL-13 (50 ng/ml) to induce myofibroblast differentiation. Additionally, M-CSF and IL-13 were refreshed in the media of mouse monocytes after 3 days of culture. Monocytes were allowed to differentiate for 5 days, and counted based on morphology as previously described [73].

Myofibroblasts were de-differentiated by addition of antibodies (α-αMβ2, α-αM, α-α3) for 7 days after the 5 day differentiation was completed. Antibodies were free of azide, glycerol, and other preservatives.

### Culture of human and mouse fibroblasts

Human fibroblasts (MRC-5, ATCC, Manassas, VA) and mouse fibroblasts (NIH-3T3, ATCC) were cultured in SFM composed for human cells, with 1x concentrations of additives. 5 ng/ml TGFβ (Peprotech) was added to induce myofibroblast formation [74]. Cells were cultured at 10,000/cm^2^, and TGFβ was refreshed in cultures weekly to maintain the myofibroblast phenotype, if necessary. Myofibroblasts were de-differentiated by addition of antibodies for 7 days.

### mRNA purification and RNAseq

mRNA was purified using the Trizol method [75]. RNA sequencing was performed by the University of Chicago Center for Research Informatics.

### Flow cytometry

Cultured myofibroblasts were lifted with ice cold trypsin-EDTA (Sigma), followed by mechanical agitation by a rubber policeman. Myofibroblasts were fixed and permeabilized using using Cytofix/Cytoperm (BD biosciences, Franklin Lakes, NJ), and live-dead stained (live-dead aqua, ThermoFisher) per manufacturer’s instructions. Antibodies used were α-collagen I (Biolegend, San Diego, CA), anti-α smooth muscle actin (αSMA) (R and D systems, Minneapolis, MN), α-ki-67 (BD biosciences), and α-talin2 (R and D systems). Compensation was performed via UltraComp beads (ThermoFisher) per the manufacturer’s instructions.

### Immunofluorescence

Human monocytes from 3 donors were culture in 8-well chamber slides (Millipore-Sigma) in SFM, and allowed to become myofibroblasts over 5 days. Myofibroblasts were then treated with antibodies against integrins α3 (α-α3), -M (α-αM), and αMβ2 (α-αMβ2) at 500 ng/ml for 1 week. After 1 week of de-differentiation, the slides were dried quickly using the airflow from a laminar flow hood, in order to preserve cellular morphology as accurately as possible. Cells were then fixed with ice cold 4% PFA, permeabilized with saponin (Sigma). Primary antibodies (α-talin2, novus) were added at 5 µg/ml overnight. Cells were gently washed 3 times in PBS, and were exposed to DAPI and F-actin-phalloidin- 488 (ThermoFisher) for 1 min. Cells were mounted using water-based mowiol mounting media (Southern Biotech, Birmingham, AL) to preserve fluorescence. Slides were imaged immediately using a Confocal microscope (Olympus, Shinjuku City, Tokyo).

### Legendplex detection by ELISA

Supernatant from cultured human and mouse myofibroblasts were thawed at 4C, and centrifuged at 4C and 10,000 xG to pellet cell debris. Supernatant was taken and added to 96 well round bottom plates, in duplicate. Legendplex (Biolegend) beads against general inflammation markers were added, according to the manufacturer’s instructions. Sample readouts were normalized to each individual donor control.

### Synthesis of peptide conjugated antibody

Conjugation of decorin’s collagen-binding peptide (CBP) (LRELHLNNNC) was performed as described previously [43]. Monoclonal antibodies, including mouse α-αM (clone ICRF44 for human experiments, clone M1/70 for mouse experiments, Biolegend), mouse anti-human/mouse α3 (α-α3, 3F9G4, Proteintech, Rosemont, IL), and mouse anti-human MAC1 (α-αMβ2, CBRM1/5, Biolegend) were incubated with 30 fold molar excess of sulfo-SMCC cross-linker (ThermoFisher) for 30 min at room temperature. The sulfo-SMCC-antibody was then mixed with 20, 30, and 40 fold molar excess CBP peptide (LRELHLNNNC) for 1 hr at room temperature, resulting in CBP-conjugated anti-integrin antibodies (CBP-α-αMβ2, CBP-α-αM, CBP-α-α3). The peptide was more than 95% purity, (Genscript, Piscataway, NJ).

### Matrix-assisted laser desorption/ionization-time-of-flight (MALDI-TOF) mass spectroscopy

MALDI-TOF was performed as previously described [43]. Briefly, MALDI-TOF was performed on CBP- conjugated anti-integrin antibodies (CBP-α-αM, CBP-α-α3, CBP-α-αMβ2) using a UltrafleXtreme MALDI TOF/TOF instrument or a Bruker AutoFlex III Smartbeam MALDI TOF. Spectra were collected using Bruker flexControl software and processed with analysis software Bruker flexAnalysis or MATLAB (MathWorks). The matrix used was a saturated solution of α-cyano-4-hydroxycinnamic acid (Sigma-Aldrich) or sinapic acid (Sigma-Aldrich), was prepared in 50:50 (v/v) acetonitrile:(1% trifluoroacetic acid in water) as a solvent. The analyte in phosphate-buffered saline (PBS) (5 μl, 0.1 mg/ml) and the matrix solution (25 μl) were then mixed, and 1 μl of that mixture was deposited on the MTP 384 ground steel target plate. The drop was allowed to dry in a nitrogen gas flow, which resulted in the formation of uniform sample/matrix coprecipitate. All samples were analyzed using the high mass linear positive mode method with 5000 laser shots at a laser intensity of 75%. The measurements were externally calibrated at three points with a mix of carbonic anhydrase, phosphorylase B, and BSA.

### In vivo biodistribution study

An in vivo biodistribution study was conducted as previously described [43], with minor adjustments.

Fluorescently labeled Cy7-antibodies (CY7-α-αMβ2, CY7-α-αM, CY7-α-α3) were conjugated using sulfo- Cy7 *N*-hydroxysuccinimide ester (Lumiprobe) according to the manufacturer’s instruction. Unreacted Cy7 was removed by dialysis against PBS.

To make fluorescently labeled Cy7-CBP-antibody, antibodies were incubated with eightfold molar excess of SM(PEG)24 cross-linker (ThermoFisher) for 30 min at room temperature. Unreacted cross-linker was removed using a Zeba spin desalting column (Thermofisher, Waltham, MA), and then 30-fold molar excess of Cy7-labeled CBP ([Cy7]LRELHLNNNC[COOH], Genscript, >95% purity) was added and reacted for 30 min at room temperature for conjugation to the thiol moiety on the C residue. Unreacted Cy7- CBP was removed by dialysis against PBS, resulting in antibodies labeled with CY7 and CBP: CY7-CBP-α- αMβ2, CY7-CBP-α-αM, and CY7-CBP-α-α3.

7 days following bleomycin insult or UUO surgery, 50 µg Cy7-CBP-antibody or Cy7-antibody were injected via tail vein. 48 hr later (in the case of lungs, Figure 3) or 24 hours later (in the case of kidneys, Figure 7), heart, lungs, spleen, kidneys, and liver were resected and imaged via IVIS (Xenogen) under the following conditions: *f*/stop, 2; optical filter excitation, 710 nm; emission, 780 nm; exposure time, 5 seconds; small binning.

### Bleomycin induced pulmonary fibrosis model

Male and female mice were acquired at 8 weeks of age (Jackson laboratories, Bar Harbor, ME) with the intent to be used at 12 weeks of age. Due to delays regarding COVID19 lockdown and the allowed resumption of non-COVID19 research, the mice were 32 weeks old when the study began. Mouse lungs were instilled with 0.075 units bleomycin (75 µg, Fresenius Kabi, Switzerland) suspended in endotoxin-free PBS, as previously described [43]. First, mice were anesthetized via isoflurane inhalation (2%). Mice were then placed upright on an angled surface, their tongue pulled to the side, and a 200 µl narrow pipet was placed at the entrance of their throat. 50 µl of bleomycin/PBS was dispensed to the entrance of the throat, and mice were allowed to inhale. Administration to the lungs was confirmed by listening to the mouse’s breathing for popping noises. Mice were then weighed and placed on a heating pad to recuperate.

Following bleomycin insult, mice were injected i.v. with 50 µg of antibody (α-αMβ2, α-αM, α-α3) or CBP- antibody (CBP-α-αMβ2, CBP-α-αM, CBP-α-α3) via tail-vein injection. The dose schedule was 7, 9, 11, 14, 16, and 18 days following bleomycin insult. Mice were euthanized at 21 days post insult via injecting of euthasol (Covetrus, Portland, ME) instead of CO2 inhalation, which could damage the lungs.

### Lung resection and fibrosis scoring

Lungs were harvested, and perfused with 5 ml of PBS via cardiac puncture. After resection, the right and left lobes were separated. The left lobe was fixed in 4% paraformaldehyde overnight, mounted in paraffin, sectioned into 5 μm slices, and stained using Masson’s trichrome. Stained lungs were scanned at high resolution using a CRi Panoramic SCAN 40x Whole Slide Scanner (Perkin-Elmer, Waltham, MA), and were read for fibrosis using a modified Ashcroft method, as previously described [56]. Lungs were read unlabeled by a researcher uninvolved with animal treatment.

The right lobe of the lung was frozen, and dehydrated using a tissue lyophilizer (Labconco, Kansas City, MO), weighed, and was assessed for collagen content by hydroxyproline assay [76]. Briefly, dried lungs were digested in 6N HCl/PBS at 100C for 24 hr. Supernatant from this digestion was added to 96 well plates and treated sequentially with chloramine-T solution and Ehrlich’s solution at 65C for 15 min to facilitate the color change reaction. Color was read at 561 nm. Quantification was provided by use of a hydroxyproline (Sigma) dilution series, which was transformed into a standard curve.

### Kidney unilateral ureteral obstruction (UUO) fibrosis model

UUO surgery was performed as previously described [77], with adjustments. Briefly, Mice were anesthetized via 2% isoflurane inhalation, and injected with meloxicam (1 mg/kg), buprenorphine (0.1 mg/kg) in a saline solution, subcutaneously. Briefly, mice were laid on their right side and an abdominal incision used to visualize the left ureter. The left ureter was ligated in the middle section of the ureter with two ties (2mm apart) using 7-0 silk sutures. Peritoneum is then closed with 5-0 vicryl and skin is closed with 5-0 nylon.

1 week following UUO ligation, the mice were injected with 50 µg of antibody (α-TGFβ, α-α3, α-αMβ2) or CBP-antibody (CBP-α-TGFβ, CBP-α-α3, CBP-α-αMβ2) via tail-vein injection. 2 weeks post UUO, and 1 week post injection, mice were sacrificed via CO2 inhalation, and their kidneys harvested. At this point, we checked to make sure that the UUO ligation was still in place, and in each case it was.

### Assessment of fibrosis in kidneys

Right (healthy) and left (fibrotic) kidneys were placed in 4% PFA for 24 hours, mounted in paraffin, sectioned into 5 mm full kidney slices, and stained using immunohistochemistry (IHC) for collagen I (1:4000, polyclonal rabbit, lifespan biosciences, Seattle WA) via a Bond-Max autostaining system (Leica biosystems, Lincolnshire, IL). Stained kidneys were scanned at high resolution using a CRi Panoramic SCAN 40x Whole Slide Scanner (Perkin-Elmer).

Images were equalized in size, and converted to .tif files using CaseViewer. Images were then imported into imageJ, scale set for conversion between microns and pixels, and deconvoluted with the “H DAB” deconvolution option. The resulting blue image was thresholded at 215 to see how many pixels were negative for collagen I, and the brown (IHC positive) image thresholded at 185 to see how many pixels were positive for collagen I. Machine-staining allowed these kidneys to be compared with high reproducibility.

### Blood analysis for markers of kidney damage

At the time of euthanasia, blood was collected via submandibular bleed into protein low-bind tubes and allowed to coagulate for 2 hours on ice. Coagulated blood was then centrifuged at 10,000 xG for 10 min, and serum collected. Serum was then diluted 4x in MilliQ water before being placed on deck on an Alfa Wassermann VetAxcel Blood Chemistry Analyzer. All tests requiring calibration were calibrated on the day of analysis and quality controls were run before analyzing samples. Serum tests were run according to kit instructions, and creatine kinase was normalized to calcium ion concentrations where indicated to account for sample hemolysis.

### Statistical analysis

Statistical analyses were performed using GraphPad Prism software, and *P* < 0.05 was considered statistically significant. Either Student’s t-test, 2-way ANOVA (with Fisher’s LSD post test), or 1-way ANOVA with Welch’s correction was used to compare groups.

## Acknowledgements

We thank the Human Tissue Resource Center of the University of Chicago for histology analysis. We thank the Integrated Light Microscopy Core of the University of Chicago for Imaging. We thank the Genomics core at the University of Chicago for RNAseq analysis. We’d also like to acknowledge blind scoring from Margo MacDonald, and guidance on fibrosis models from Anne Sperling.

## Abbreviations

alpha-smooth muscle actin: (αSMA)
CBP-functionalized blocking antibodies against αMβ2: (CBP-α-αMβ2)
CBP-functionalized blocking antibodies against αM: (CBP-α-αM)
CBP-functionalized blocking antibodies against α3: (CBP-α-α3)
collagen-binding peptide: (CBP)(LRELHLNNNC)
fetal calf serum: (FCS)
Fibrillar adhesions: (FBs)
Fluorescently labeled Cy7-blocking antibodies against α3: (CY7-α-α3)
Fuorescently labeled Cy7-blocking antibodies against αMβ2: (CY7-α-αMβ2)
Fluorescently labeled Cy7-blocking antibodies against αM: (CY7-α-αM)
focal adhesions: (FAs)
Gi14: (antibody which stabilizes the association of integrin α2β1)
Human fibroblasts (MRC-5) heterodimer αMβ2: (called MAC1)
Idiopathic pulmonary fibrosis: (IPF)
IL-12 subunit p40: (IL12p40)
Integrin-α3: (α3)
Integrin-αM: (αM)
Integrin-β2 (β2): (CD18)
macrophage-chemotactic protein-1: (MCP1)
mouse fibroblasts: (NIH-3T3)
blocking antibodies against αMβ2: (α-αMβ2)
blocking antibodies against αM: (α-αM)
blocking antibodies against α3: (α-α3)
peripheral blood mononuclear cells: (PBMC)
phosphate buffered saline: (PBS)
serum-free media: (SFM)
Transforming growth factor β: (TGFβ)
Tumor necrosis factor: (TNFα)
Unilateral ureteral obstruction: (UUO)

## Author’s contributions

Conceptualization: MJVW Methodology: MJVW

Investigation: MJVW, MO, JEGM, KK, MMR, ATA Visualization: MJVW

Funding acquisition: JAH

Project administration: MJVW, JAH Supervision: MJVW, JAH

Writing – original draft: MJVW

Writing – review & editing: MJVW, JAH

## Ethics approval

All the animal experiments performed in this work were approved by the Institutional Animal Care and Use Committee of the University of Chicago.

## Supplemental Figures

**Supplemental figure 1:**
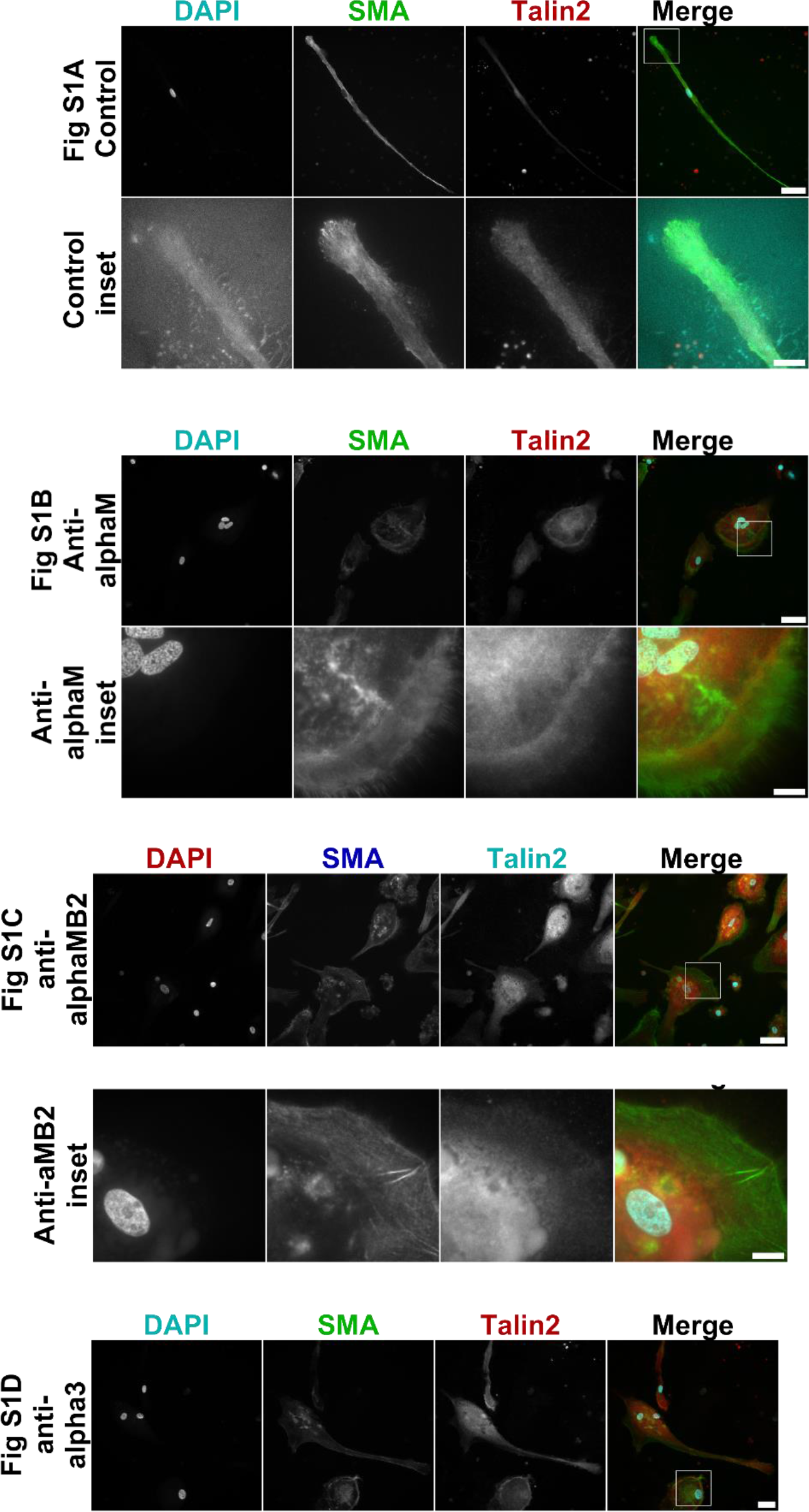
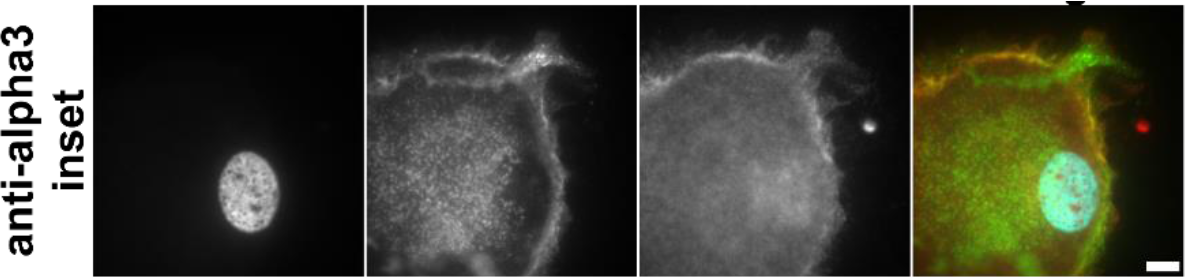
Antibodies against integrins α3, αM, and αMβ2 de-differentiate myofibroblasts, and remove focal adhesion subcellular structures (FAs). Human monocyte-derived myofibroblasts were treated with 500 ng/ml α-αM, α-αMβ2, α-α3. (A) Untreated population of myofibroblasts, inset on nucleus and focal adhesion at the tip of the myofibroblast. (B) Myofibroblasts treated with α-αM, inset on edge of de-differentiated myofibroblast. Actin staining is diffuse. (C) Myofibroblasts treated with α-αMβ2, inset on the edge of de-differentiated myofibroblast. Actin staining is diffuse. (D) Myofibroblasts treated with α-α3, inset on the edge of de-differentiated myofibroblast. Actin staining is diffuse. Red = talin2, green = αSMA, blue = DAPI. Size bar = 30 μm, inset size bar = 10 μm.

**Supplemental figure 2:**
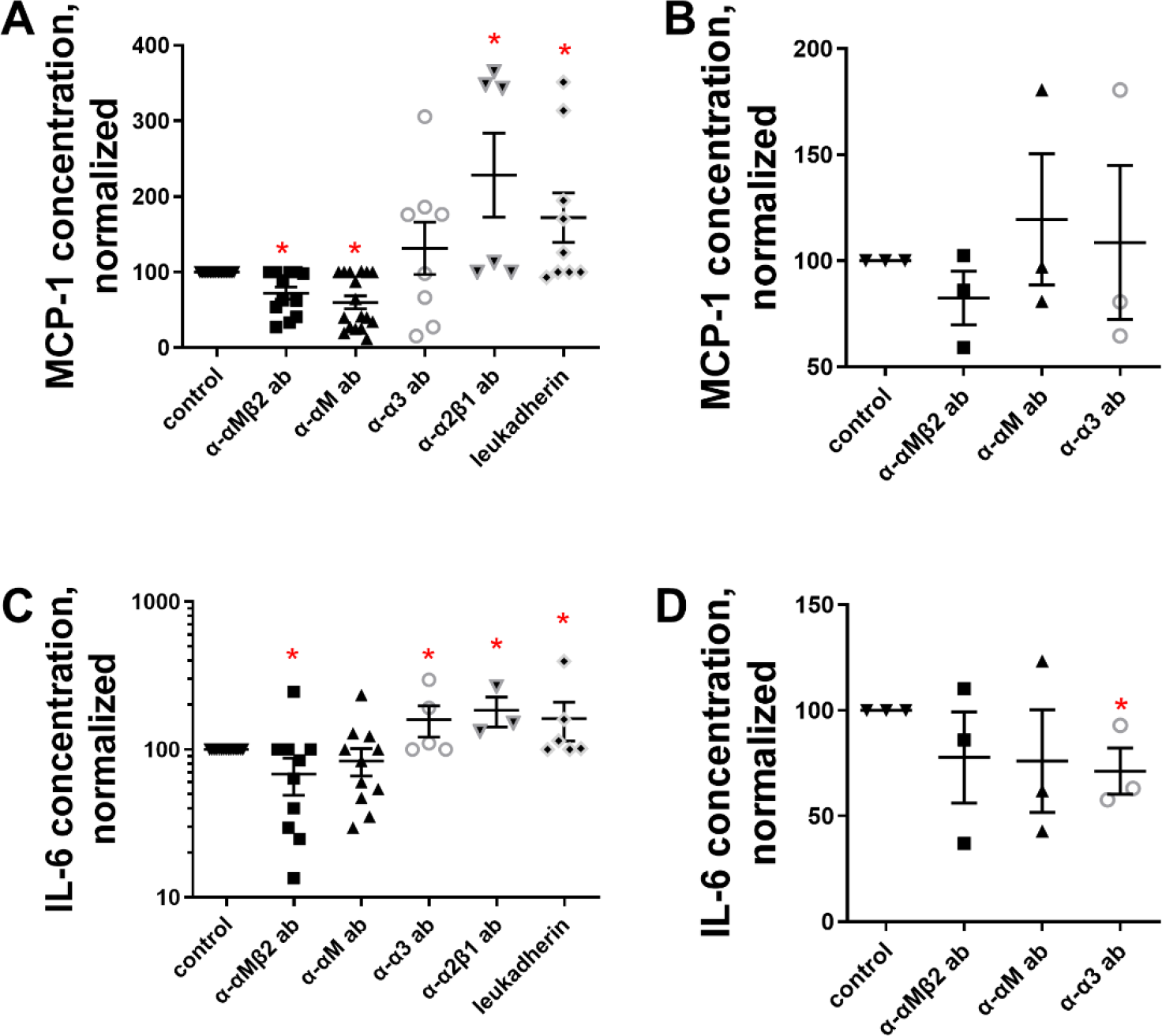
Antibodies against integrins α3, αM, and αMβ2 reduce the pro-fibrotic secretome of human myofibroblasts. Human monocyte-derived myofibroblasts and fibroblast-derived myofibroblasts were treated with 500 ng/ml of the indicated antibodies (or 2 ng/ml leukadherin), and the conditioned media after 1 week was assessed via Legendplex ELISA. Secreted proteins over the detection limit were: (A) MCP-1 from monocyte-derived myofibroblasts, (B) MCP-1 from fibroblast- myofibroblasts. (C) IL-6 from monocyte-derived myofibroblasts, IL-6 from fibroblast-derived myofibroblasts. n ranges from 3 to 15. * = statistical significance of P < 0.05, < 0.01, or < 0.001, significance vs control, Student’s t-test.

**Supplemental figure 3:**
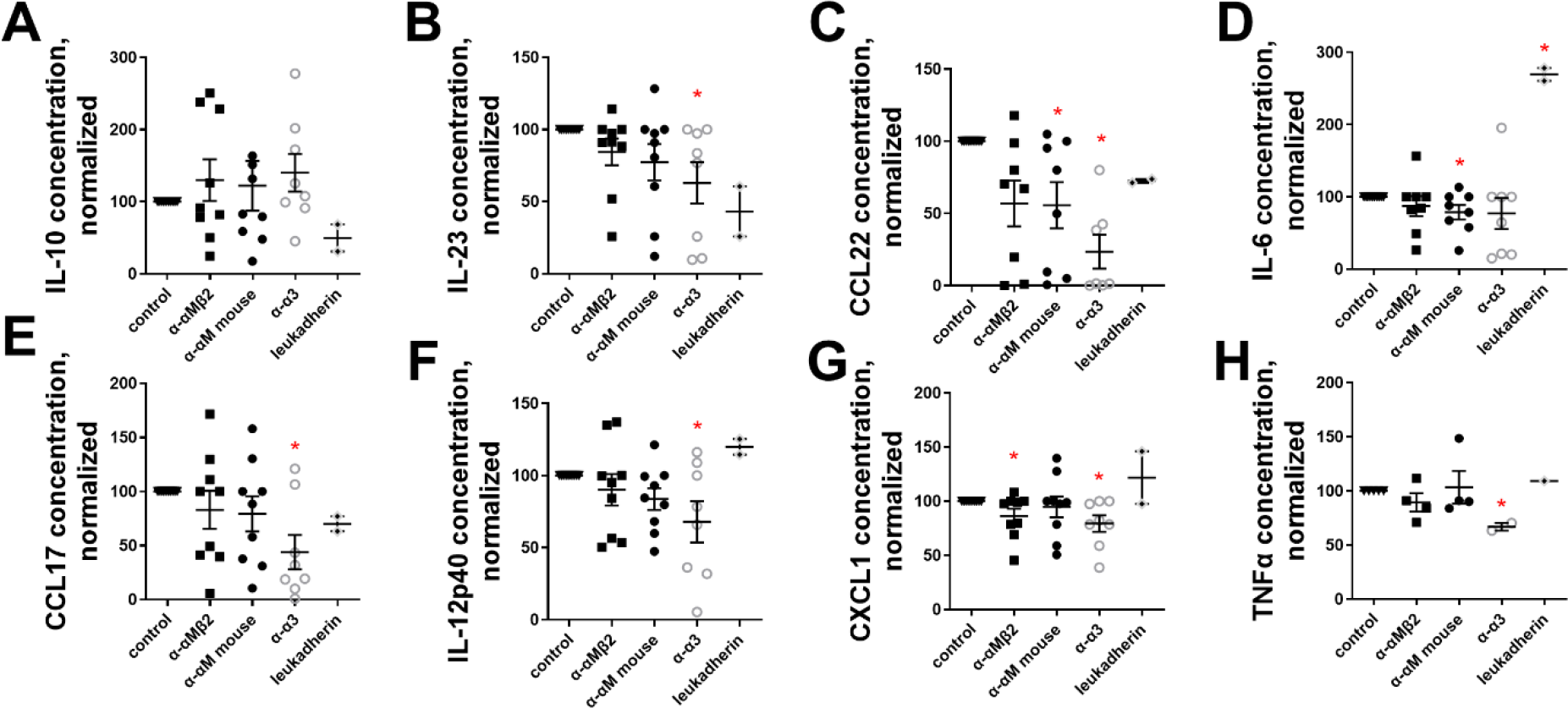
Antibodies against integrins α3, αM, and αMβ2 reduce the pro-fibrotic secretome of mouse myofibroblasts. Mouse monocyte-derived myofibroblasts were treated with 500 ng/ml of α-α3, α-αM, or α-αMβ2, and the conditioned media after 1 week was assessed via Legendplex ELISA. Secreted proteins over the detection limit were: (A) IL-10, (B) IL-23, (C) CCL22, (D) IL-6, (E) CCL17, (F) IL-12 subunit p40, (G) CXCL1, (H) TNF-α. n ranges from 2 to 8. * = statistical significance of P < 0.05, < 0.01, or < 0.001, significance vs control unless otherwise indicated, Student’s t-test.

**Supplemental figure 4:**
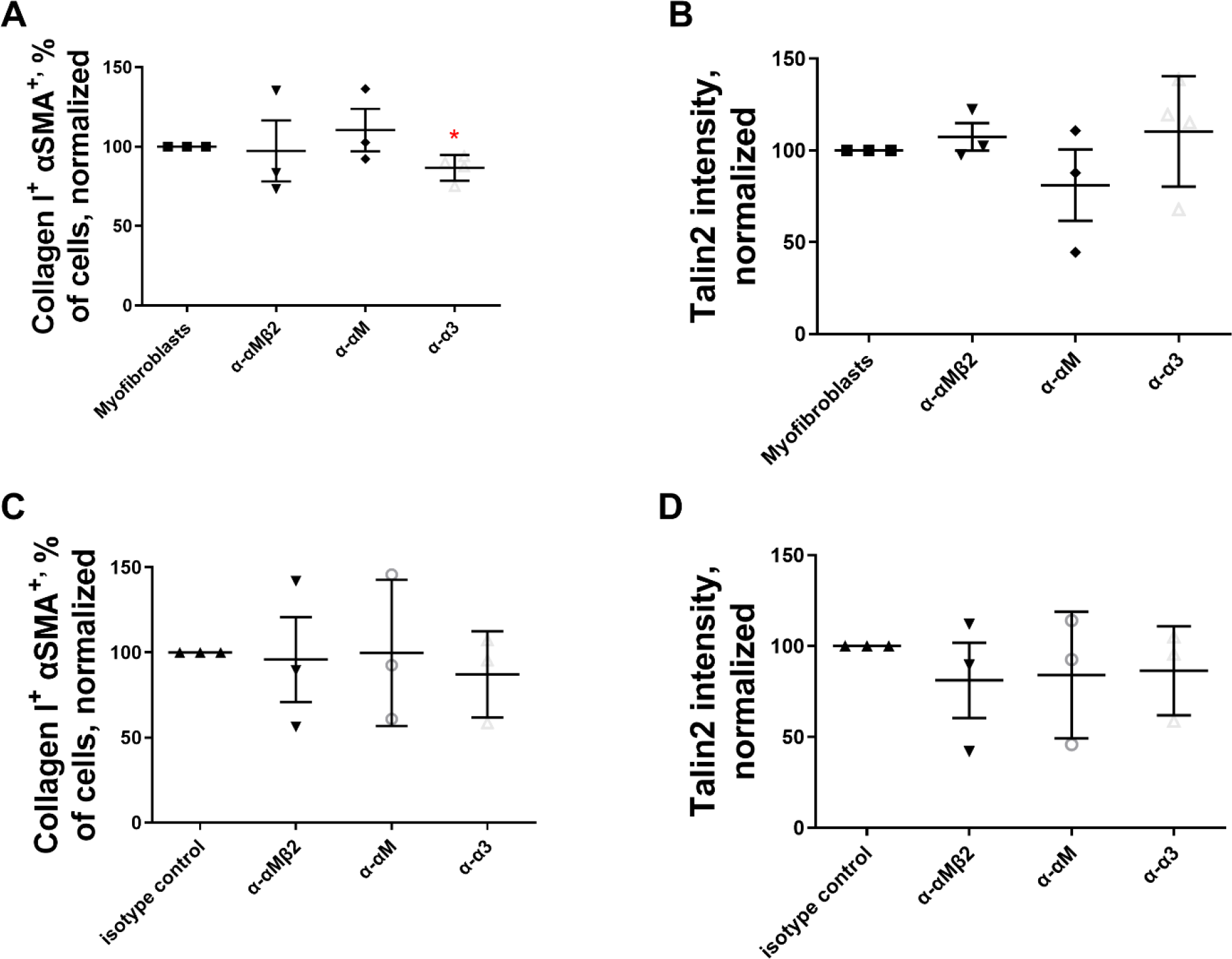
α-α3 de-differentiates human fibroblast-derived myofibroblasts, while α-αM, and α-αMβ2 do not. Human (A and B) and mouse (C and D) fibroblasts were induced to become myofibroblasts with 5 ng/m TGFβ, treated for 1 week with the indicated antibodies at 500 ng/ml, and analyzed via flow cytometry for (A and C) αSMA^+^ and collagen I^+^ double positive cells and (B and D) talin2^+^. n ranges from 2 to 3. * = statistical significance of P < 0.05, < 0.01, or < 0.001, significance vs isotype control antibody, Student’s t-test.

**Supplemental figure 5:**
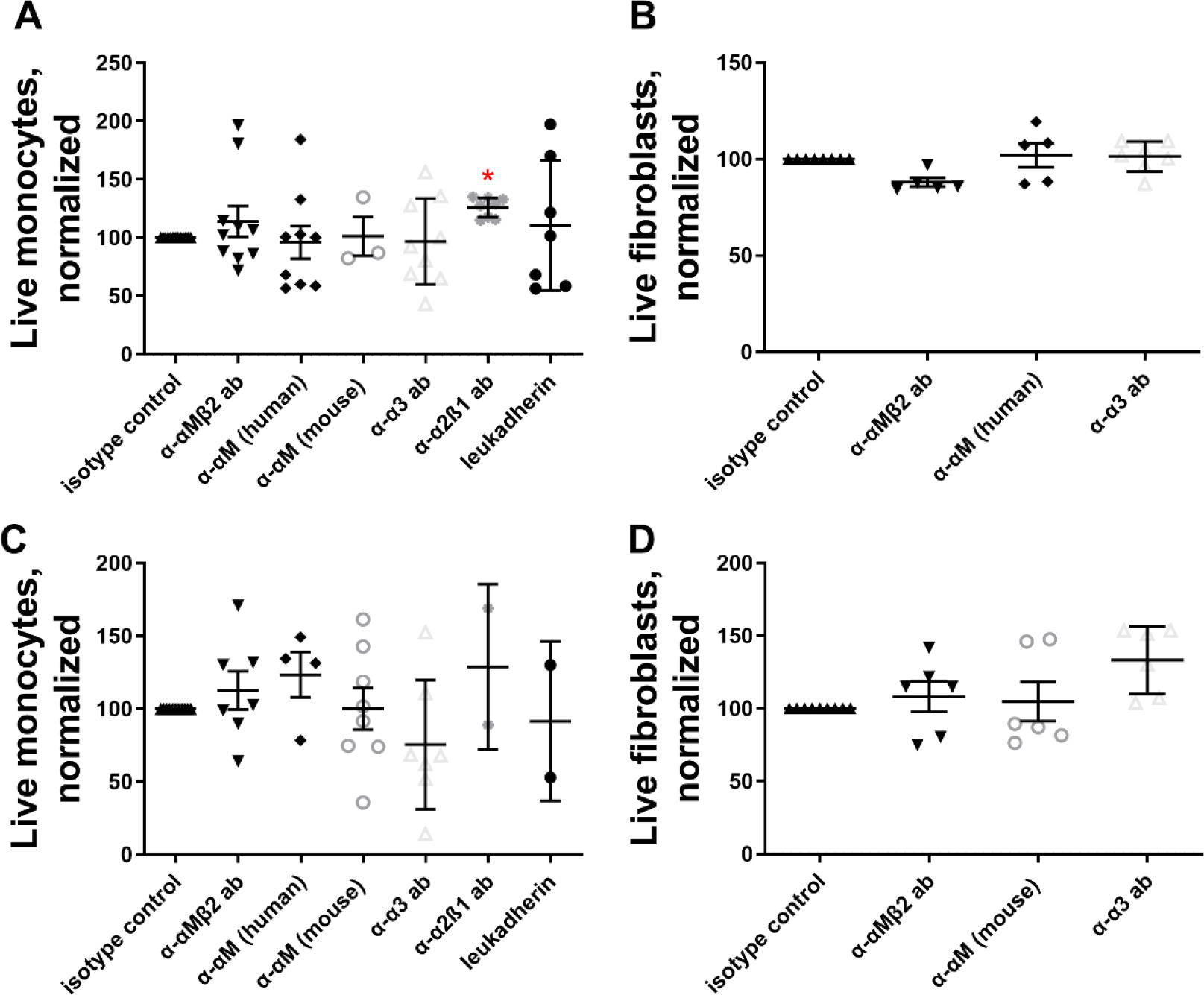
Antibodies against integrins α-α3, α-αM, and α-αMβ2 do not cause cell death. 500 ng/ml of the indicated antibodies (or 2 ng/ml leukadherin) were added to myofibroblasts for 1 week, then cells were assessed for living by live-dead aqua stain. (A) Human monocyte-derived myofibroblasts, (B) human fibroblast-derived myofibroblasts, (C) mouse monocyte-derived myofibroblasts, (D) mouse fibroblast-derived myofibroblasts. * = statistical significance of P < 0.05, < 0.01, or < 0.001, significance vs isotype control antibody, Student’s t-test.

**Supplemental figure 6:**
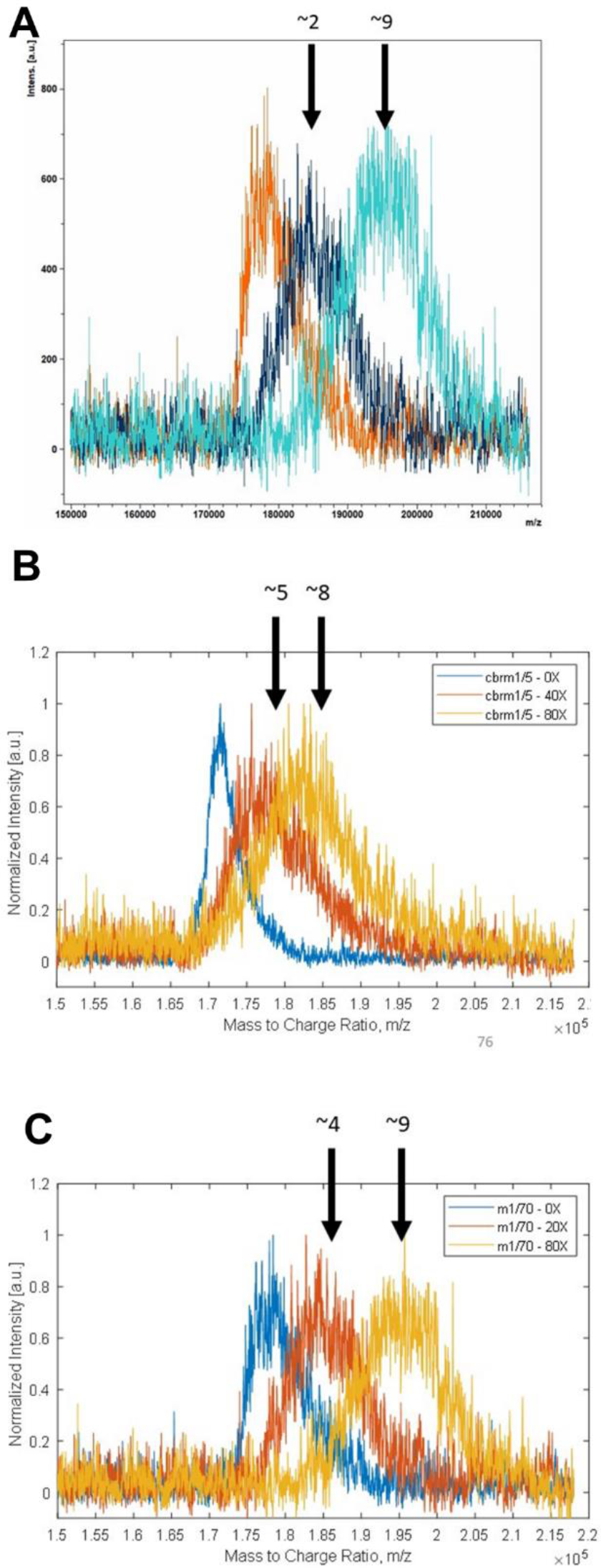
Antibodies against integrins α3, αM, and αMβ2 were conjugated with CBP at approx. 5 peptide copies per antibody molecule. Unlabeled CBP was conjugated to antibodies, and mass change measured via MALDI-TOF: (A) α-α3, (B) α-αM, (C) α-αMβ2. Since the combined molecular weight of the linker and CBP peptide is 2.5 kDa, total numbers of attached CBP peptides were derived.

**Supplemental Figure 7:**
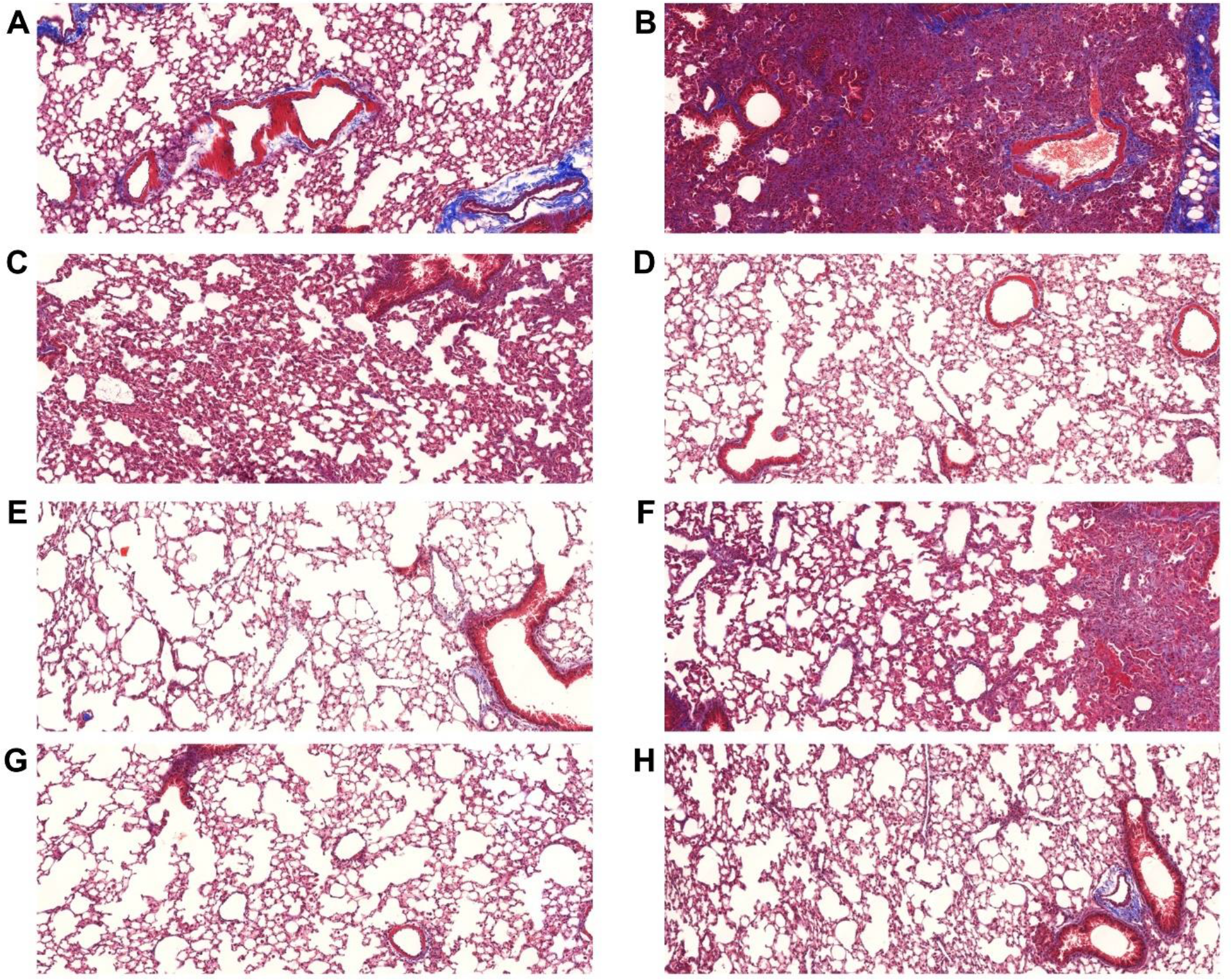
Representative images of left, single-lobed lungs stained with Massons’s trichrome. (A) Uninsulted lungs instilled with PBS. Lungs insulted with (B-H) 75 μg bleomycin, and instilled with (C) α-α3, (D) CBP-α-α3, (E) α-αM, (F) CBP-α-αM, (G) α-αMβ2, (F) CBP-α-αMβ2. Lungs were harvested 3 weeks after insult and stained via Masson’s trichrome. Scores in Figures 4, 6, and full lungs in figure 5.

**Supplemental Figure 8:**
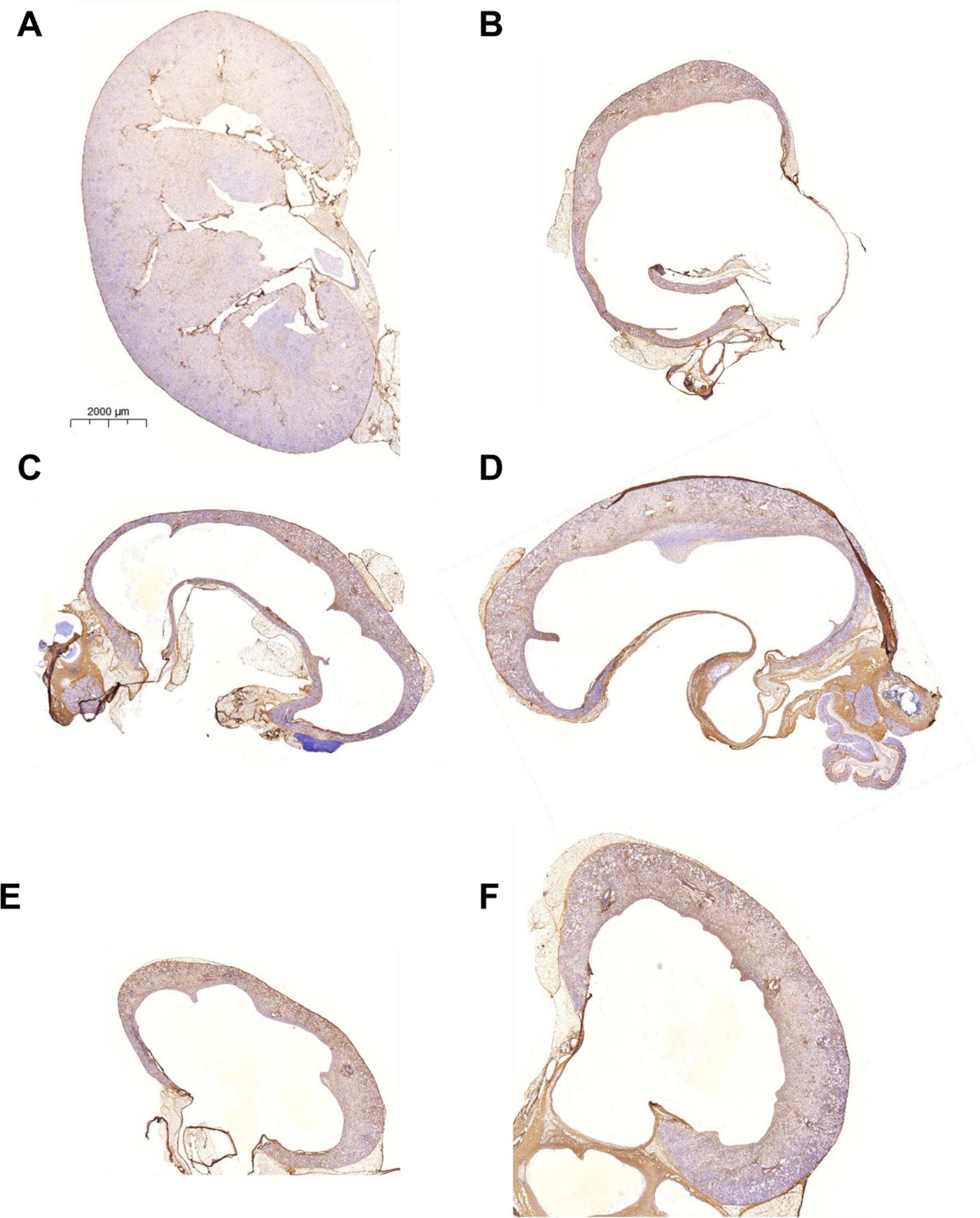
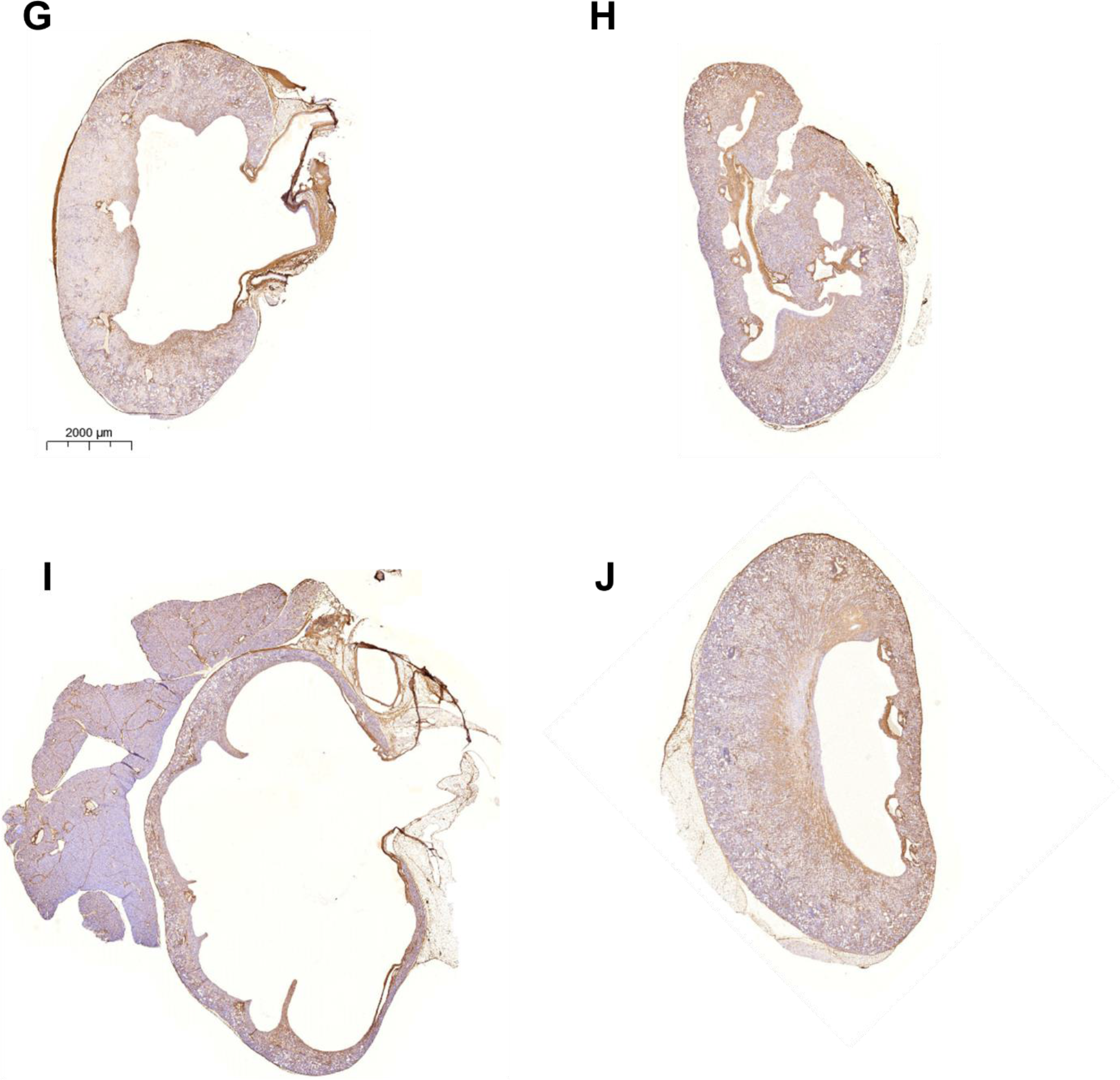
Representative images of kidneys, immunohistochemistry stained for collagen I. (A) Uninsulted kidney, (B) untreated UUO kidney after 2 weeks, UUO kidney (2 weeks after UUO, 1 week after antibody injection) treated with 50 micrograms of (C) isotype control antibody, (D) CBP- isotype control antibody, (E) α-TGFβ, (F) CBP-α-TGFβ, (G) α-α3, (H) CBP-α-α3, (I) α-αMβ2, (J) CBP-α- αMβ2. Data in Figure 7.

**Supplemental Figure 9:**
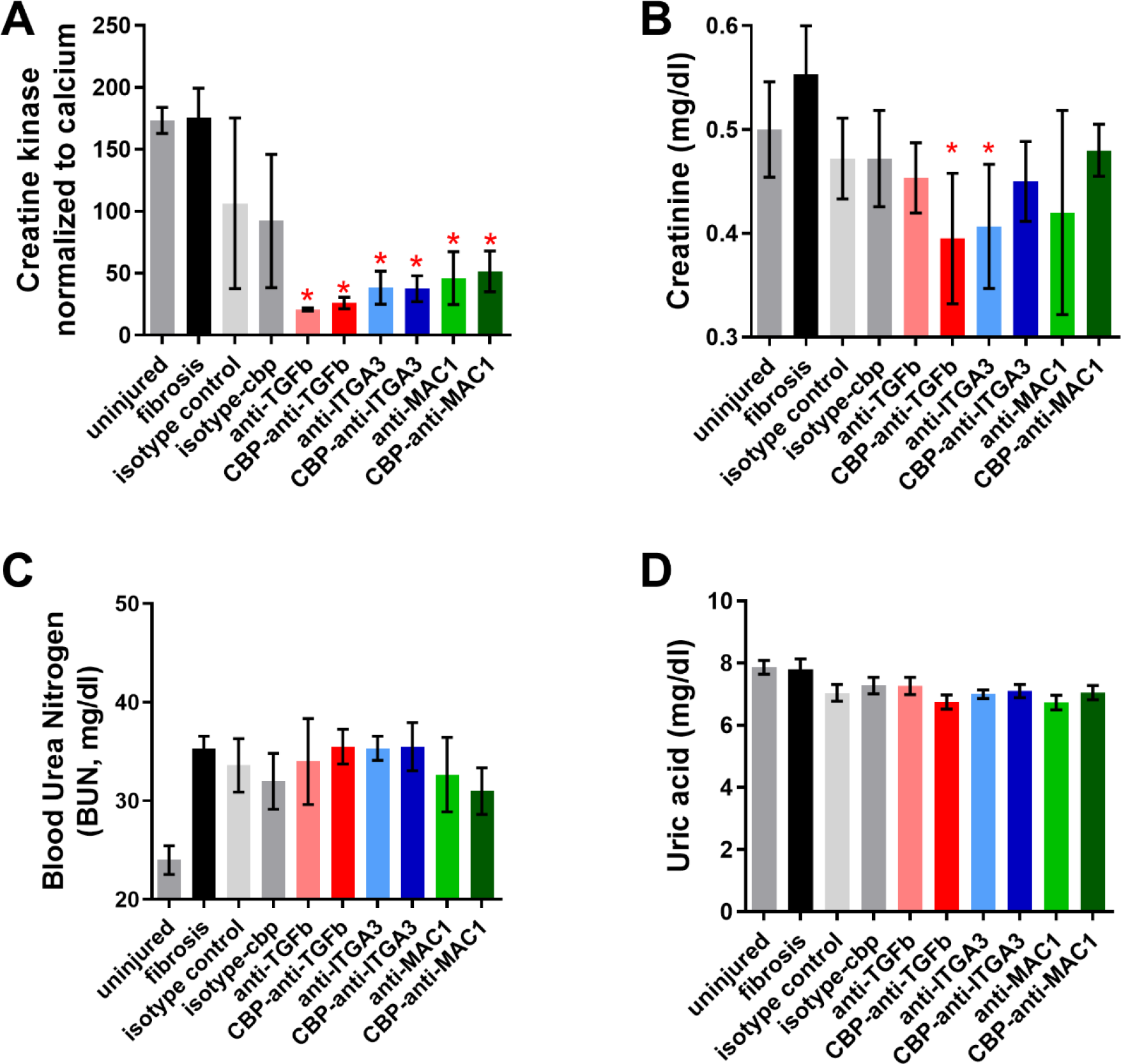
Blood analysis from kidney UUO model. Blood was taken from mice directly before sacrifice at 2 weeks post UUO, 1 week post antibody injection. Concentrations in serum of (A) creatine kinase, (B) creatinine, (C) blood urea nitrogen, and (D) uric acid. * = statistical significance of P < 0.05, < 0.01, or < 0.001, significance vs fibrosis, one way ANOVA, no post test.

**Supplemental Figure 10:**
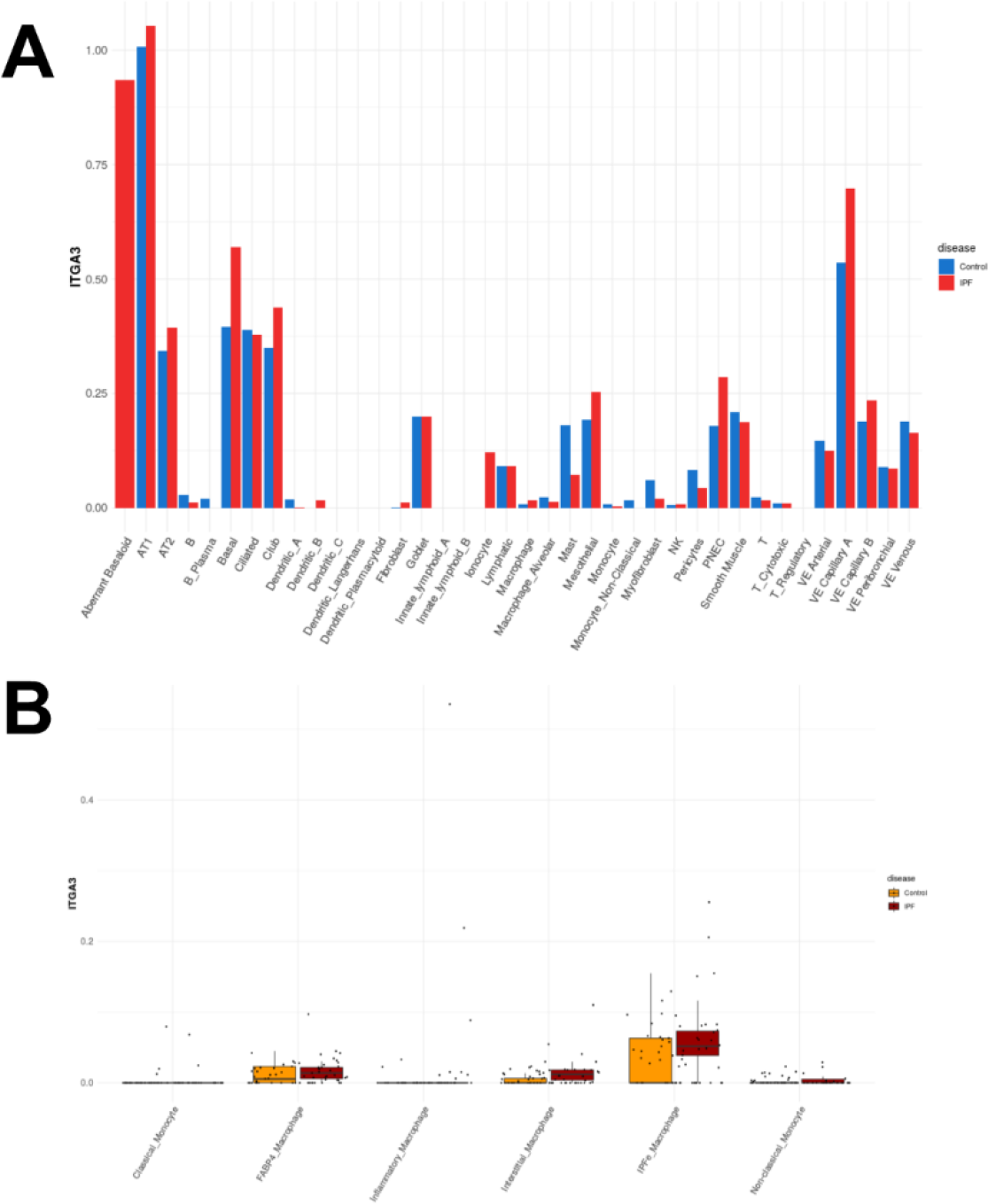
Integrin α3 is upregulated among monocyte-derived cell types found in IPF. (A) Integrin α3 transcript frequency in cells found in IPF. (B) Integrin α3 transcript frequency by individual patient for selected immune cell subtypes. All data and figures from the IPF Atlas, Kraminski/Rosas dataset.

**Supplemental Figure 11:**
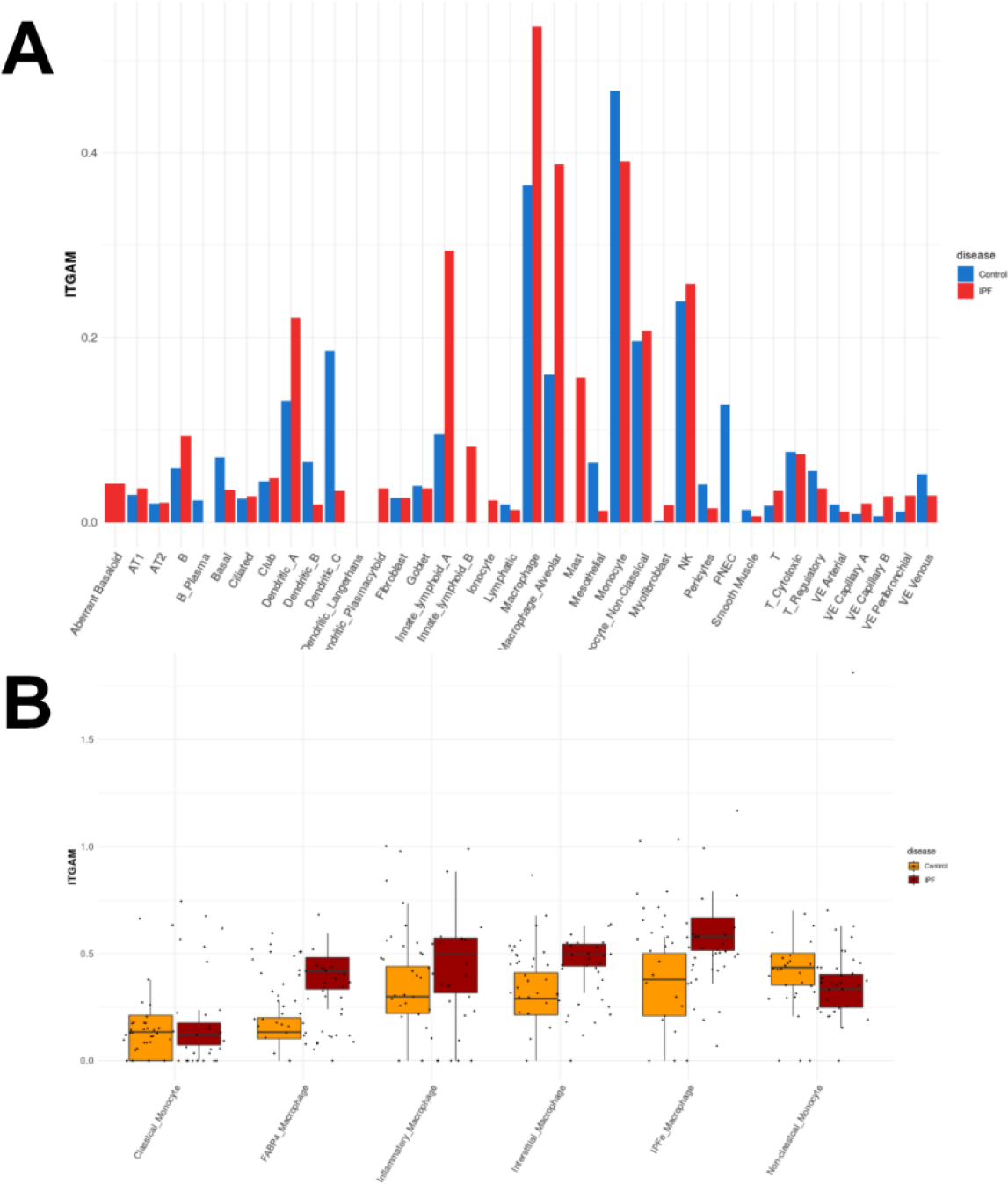
Integrin αM is upregulated among monocyte-derived cell types found in IPF. (A) Integrin αM transcript frequency in cells found in IPF. (B) Integrin αM transcript frequency by individual patient for selected immune cell subtypes. All data and figures from the IPF Atlas, Kraminski/Rosas dataset.

**Supplemental Figure 12:**
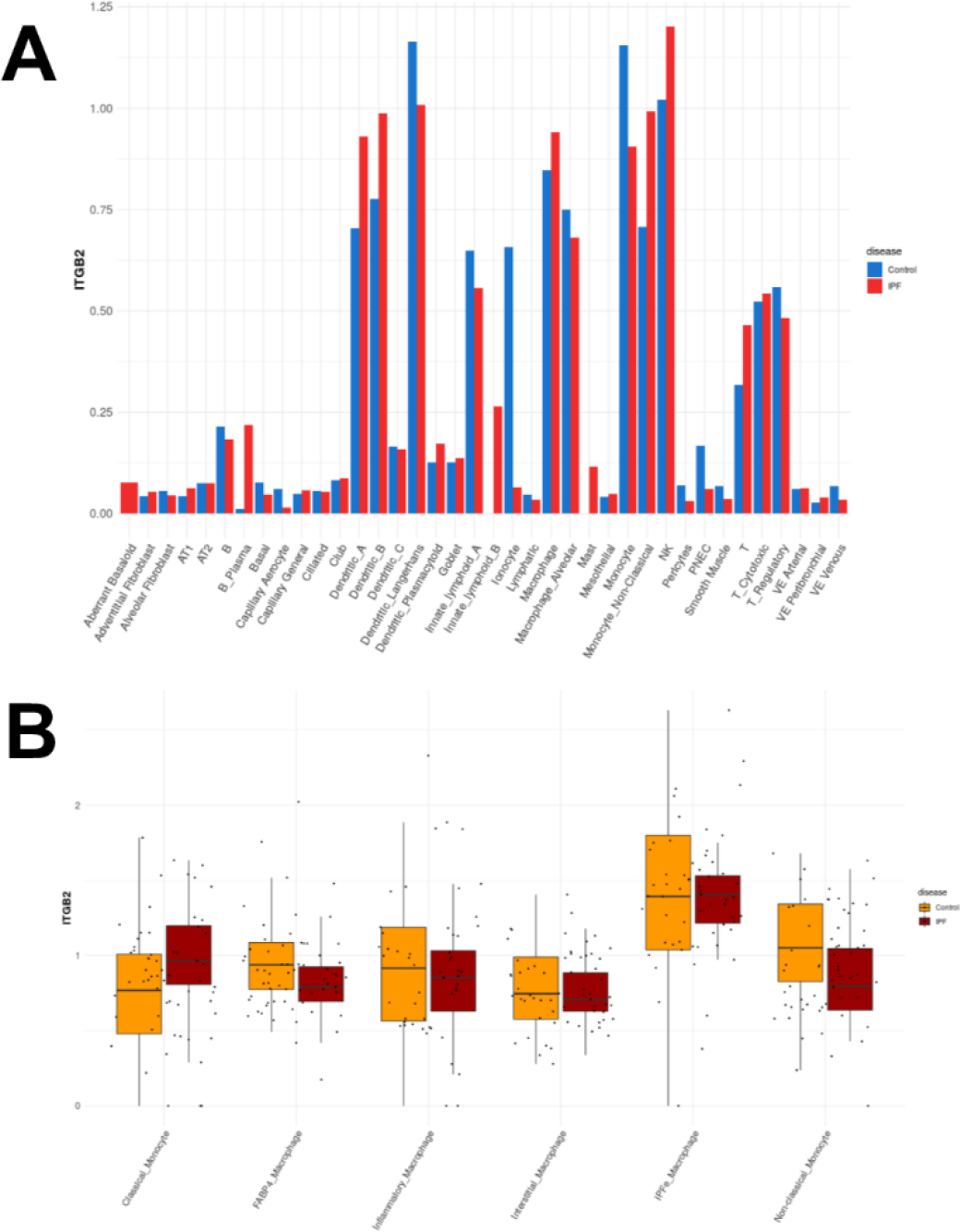
Integrin β2 is upregulated among monocyte-derived cell types found in IPF. (A) Integrin β2 transcript frequency in cells found in IPF. (B) Integrin β2 transcript frequency by individual patient for selected immune cell subtypes. All data and figures from the IPF Atlas, Kraminski/Rosas dataset.

